# Protective and pathogenic antibody responses from a primate *Shigella* outbreak inform vaccine design

**DOI:** 10.64898/2025.12.16.694763

**Authors:** Robert M. Gallant, Paul Savarino, Sophia Pulido, Morgan Gilman, Ti Lu, Jennifer Hayes, Nicholas L. Xerri, Torrey Williams, Mark A. Ochoa, Zackary Dietz, Eric Peterson, Timothy Scott, Faye Hartmann, Stephen G. Baker, Robert W. Kaminski, Devin Sok, Andrew C. Kruse, Wendy Picking, Saverio V. Capuano, Hayden R. Schmidt

## Abstract

There is currently no approved vaccine for *Shigella* spp., a leading cause of diarrhea that are increasingly resistant to antimicrobials. *Shigella* vaccine development is complicated in part by an incomplete understanding of the structural and molecular determinants of immunity. To address this, we isolated monoclonal antibodies (mAbs) against candidate *Shigella* vaccine antigens using samples from a *Shigella flexneri* outbreak in a non-human primate (NHP) research facility. We found that antibodies targeting the *Shigella* O-antigen (O-Ag) can undergo significant affinity maturation (>10%) to acquire broad cross-reactivity across *S. flexneri* serotypes. We also found that the virulence-associated type III secretion system (T3SS) proteins IpaD and IpaB elicit moderate T cell and robust antibody responses. T3SS antibodies either inhibit, or – surprisingly – enhance bacterial virulence *in vitro* and *in vivo* depending on their epitope specificity. These findings provide key insights into protective and deleterious immune responses against *Shigella* that directly inform vaccine immunogen design.

## Introduction

*Shigella* infection is a leading cause of diarrheal disease, accounting for over 200,000,000 cases globally each year.^1^ These cases result in over 200,000 deaths and substantial morbidity among young children in the form of linear growth faltering, stunting, and environmental enteric dysfunction^2^. As rates of resistance to fluoroquinolones and other antimicrobials increase among *Shigella* isolates,^3–6^ an effective vaccine is increasingly important. Several *Shigella* vaccine candidates are in the pipeline at various stages of development,^7^ but *Shigella* vaccines face significant hurdles to clinical adoption. Given that immunity against *Shigella* spp. tends to only protect against a narrow subset of the many clinically relevant serotypes,^8,9^ the bacteria are highly virulent (the infectious dose can be as low as 10-200 colony forming units^10^ (CFU)), and *Shigella* overwhelmingly impacts resource-constrained settings,^11^ vaccines must meet challenging standards with respect to breadth, potency, and cost of goods to be adopted.^12^

One method for maximizing vaccine efficacy is to apply rational antigen design principles to *Shigella* vaccine development (a strategy referred to as “Reverse Vaccinology 2.0”).^13^ These approaches have been highly effective for the design of vaccines against otherwise intractable pathogens of global health interest, including respiratory syncytial virus (RSV)^14^ and SARS-CoV-2,^15,16^ and show promise for the antigenically diverse human immunodeficiency virus (HIV).^17^ However, rational design of vaccine antigens requires a detailed molecular and structural understanding of both the antigen and the protective host immune response that is currently lacking for key *Shigella* antigens. While there is strong evidence that immune responses against O-antigen (O-Ag; the polysaccharide component of lipopolysaccharide that determines *Shigella* serotype^18–24)^ and type III secretion system (T3SS) proteins such as IpaB, IpaC, or IpaD^25–28^ are relevant for protection against disease, the precise components of the adaptive immune response that mechanistically drive protection have not been identified.

In December 2022, the Wisconsin National Primate Research Center (WNPRC) began experiencing a high incidence of *Shigella* infection among their non-human primates (NHPs), with over 150 animals experiencing symptomatic infection over a two-year period. While the scale of this outbreak is unusual, its occurrence is not – NHPs are natural hosts for *Shigella*, and outbreaks in zoos^29,30^ and research facilities^31–33^ are well-documented. This outbreak provided a unique opportunity to study the immune response to *Shigella* infection under more natural exposure conditions. Here, we integrated genomic and immunologic methods to characterize both the pathogen and the host immune response. We found that the outbreak was caused by two circulating lineages of *Shigella flexneri* that resembled contemporary isolates responsible for human disease. Infection elicited innate cytokine activation, antigen-specific T cell responses, and polyclonal antibody responses to key antigens. We isolated over 100 monoclonal antibodies (mAbs) against O-Ag, IpaB, and IpaD, and showed that mAbs against a given antigen exhibit epitope-dependent differences in their functional breadth and potency, with some subsets of mAbs even increasing virulence. Altogether, the insights gained from studying this outbreak in an NHP facility constitute an important step towards understanding which elements of an anti-*Shigella* immune response may be protective, with clear implications for the rational design of *Shigella* vaccine antigens.

## Results

### The WNPRC outbreak is driven by two genetically distinct clusters of *S. flexneri*

From December 2022 to January 2025, there were 169 cases of microbiologically confirmed shigellosis in the primate colony (Figure 1A). We performed classical serotyping on 151 isolates and whole-genome sequencing on 95 representative isolates to assess the phylogenetic diversity of the outbreak. Two main clusters of *Shigella flexneri* were identified: one predominantly serotype Y and the other predominantly serotype 3a/3b (Figure 1B, Supplemental Figure 1A). The serotype Y cluster was detected across four colony buildings while the serotype 3a/3b cluster was detected across three buildings, indicating substantial inter-building spread (Figure 1B). These isolates were closely related to human *S. flexneri* isolates from a 2019-2021 outbreak in Seattle^34^, and more distantly related to the *S. flexneri* 301 reference strain from Beijing, China, 1984^35^ (Figure 1B, Supplemental Figure 1A). Additionally, some staff members at the primate facility developed shigellosis possibly due to exposure to the infected animals in the NHP colony (personal communication), underscoring the clinical relevance of these isolates. Of the 90 sequenced isolates collected from NHPs tested for antimicrobial susceptibility, 57 (63%) were multidrug resistant (MDR), being resistant to ampicillin, amoxicillin, cephalexin, chloramphenicol, and doxycycline (Supplemental Figure 1B). These MDR isolates were almost entirely found within the primarily serotype Y cluster, and predominantly carried the resistance genes *bla*_OXA-1_, *catA1*, *tet(B)*, and *aadA1* (Supplemental Figure 1B).

**Figure 1:**
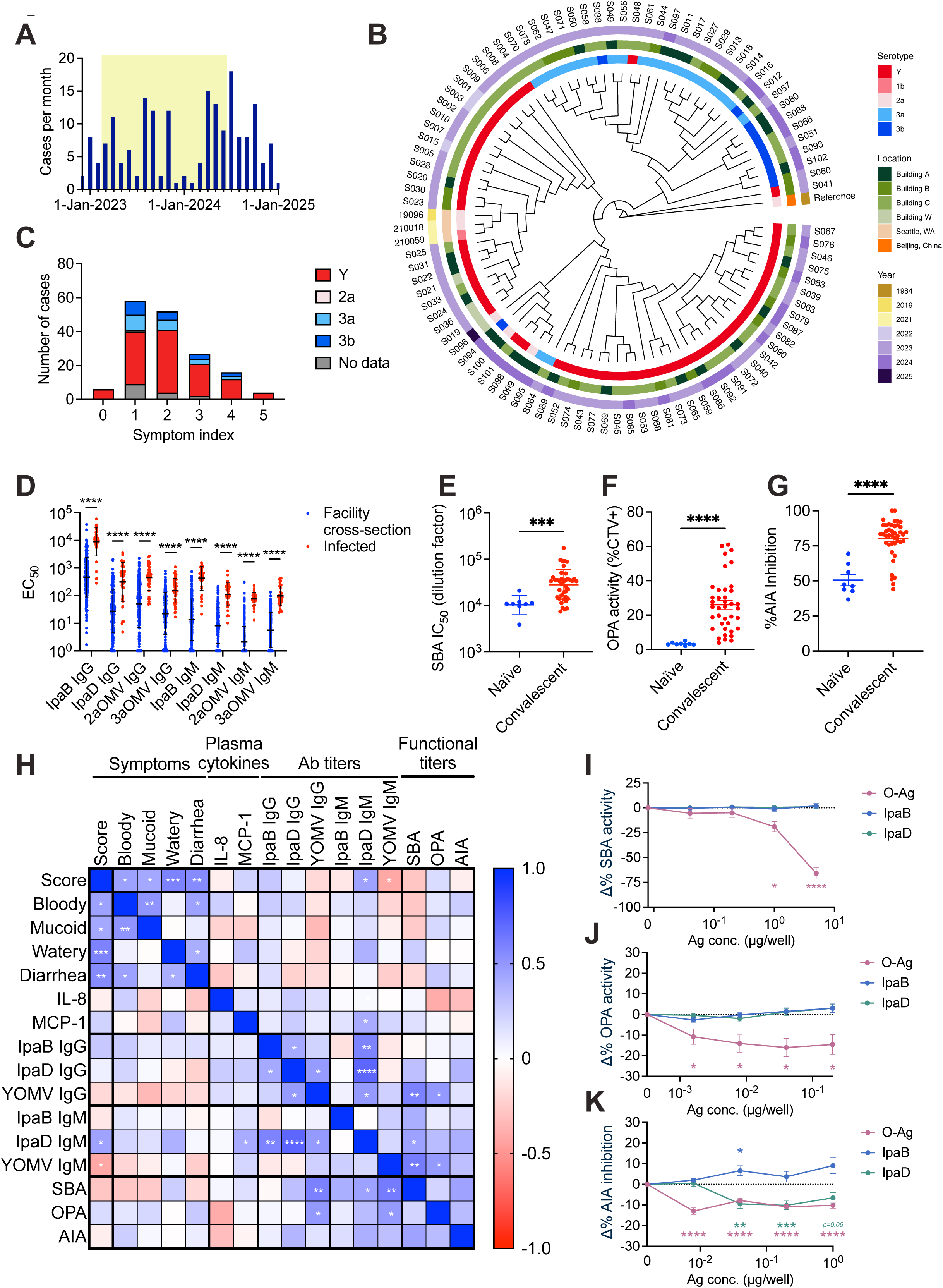
Genetic, clinical, and serological features of a *Shigella flexneri* outbreak in a NHP research facility. (A) Timeline of *Shigella* cases in the NHP research facility. Yellow shading indicates the sampling period for this study. (B) MaxPhylogenetic cladogram of *S. flexneri* isolates from the outbreak, recent human cases (19096, 210018, & 210059),^34^ and the reference strain 301.^35^ This tree is shown as a cladogram with branch lengths not proportional to evolutionary distance. Metadata (serotype, location, year) are shown circumferentially. (C) Distribution of symptom severity by serotype. 0 indicates asymptomatic infection, 1-4 indicates increasing severity, and 5 indicates euthanasia was elected or the animal was found dead. (D) IgG and IgM ELISA EC_50_ values against IpaB, IpaD, 2a OMVs, and 3a OMVs of convalescent serum collected two-weeks post detection of infection (n = 41) and an asymptomatic observational cross-section of the research facility (n = 213). Samples without measurable EC_50_ were assigned a value of 1. Geometric mean ± geometric SD, multiple lognormal Welch’s t-tests with Holm-Šídák correction. Conducted in duplicate. (E-G) Functional titers of naïve (n = 8) and two-weeks post detection of infection serotype Y-convalescent plasma (n=39). (E) Serum bactericidal assay (SBA) IC_50_, (F) opsonophagocytosis (% CellTrace Violet^+^), and (G) adhesion/invasion inhibition (% Inhibition). SBA geometric mean ± geometric SD, lognormal Welch’s t-test. OPA and AIA mean ± SEM, Welch’s t-tests. Conducted in duplicate. (H) Pearson correlation matrix of symptoms, plasma cytokines, serum antibody titers, and serum functional titers of samples collected two-weeks post detection of infection with serotype Y. Plasma cytokines, serum antibody titers, and SBA IC_50_ values are log-transformed. (I-K) Change in (I) SBA (n=10), (J) OPA (n=14), and (K) AIA (n=17) of two-weeks post detection of infection serotype Y-convalescent serum following pre-incubation with Y O-antigen (O-Ag), IpaD, or IpaB. Mean ± SEM, ordinary one-way ANOVAs with Dunnett’s multiple comparisons tests against each unblocked serum. Conducted in duplicate. \**p*< 0.05, \*\**p*< 0.01, \*\*\**p*< 0.001, and \*\*\*\**p*< 0.0001.

Regardless of infecting serotype, most NHP cases were associated with mild symptoms (a symptom index score of 1 or 2), while a minority of infections were asymptomatic or caused severe disease (Figure 1C, Supplemental Figure 2A). Symptom severity did not differ by sex (Supplemental Figure 2B), though there was a modest positive correlation with age (Supplemental Figure 2C). Additionally, infants accounted for only 12% of cases but 50% of fatal outcomes, suggesting increased susceptibility to severe disease in young primates, consistent with human epidemiological data^36^ (Supplemental Figure 2C).

### *S. flexneri* infection elicits innate and humoral immune responses in NHPs

We collected serum, plasma, and peripheral blood mononuclear cells (PBMCs) from 41 infected animals at one, two, and four-to-six weeks post detection of infection. We first measured plasma cytokines one week post detection of infection. Infected primates showed mild increases in IL-8 and MCP-1 (Supplemental Figure 2D), which is consistent with high secretion of IL-8 and MCP-1 following *Shigella* infection in various model organisms^37–39^. We observed no significant changes in other cytokines compared to naïve controls (Supplemental Figure 2D). We next compared IgG and IgM serum titers two weeks post detection of infection against IpaB, IpaD, and outer membrane vesicles (OMVs) from serotypes 2a and 3a, relative to a cross-sectional cohort of 213 asymptomatic NHPs (of unknown *Shigella* infection status) across the facility. Eight serum samples were considered polyreactive and thus excluded from antibody titer analyses due to high binding to the nonspecific control protein ovalbumin (Supplemental Figure 2E). Titers against all antigens were significantly elevated in infected animals (Figure 1D). Notably, there were several asymptomatic NHPs with antibody titers above the median infected titers, suggesting there was an undetected transmission amongst NHP colony members (Figure 1D). Time-course analysis revealed stable, elevated IgG titers by one week post detection of infection, whereas IgM responses remained lower throughout the infection, consistent with recall of pre-existing memory responses (Supplemental Figure 2F-H).

Functional antibody responses were also elevated at two weeks post detection of infection in serotype Y-infected NHPs compared to naïve controls. Convalescent plasma exhibited greater complement-mediated killing in serum bactericidal assays (SBA), greater opsonization activity in a complement-independent monocyte opsonophagocytosis assay (OPA), and greater inhibition of virulence as measured by a HeLa cell-based adhesion and invasion assay (AIA) (Figure 1E-G, Supplemental Figure 2I-K). We next evaluated whether any of these immune parameters correlated with symptom severity. IpaD-specific IgM titers were positively correlated with symptom severity, while serotype Y OMV IgM titers were inversely correlated – no other statistically significant correlations emerged (Figure 1H).

Building on these correlational findings with mechanistic experiments, we conducted antigen-specific serum blocking studies. Convalescent sera were pre-incubated with antigen to block corresponding antibodies, then tested for loss of effector function. SBA and OPA activity were significantly reduced when sera from Y-infected primates were blocked with Y O-Ag (Figure 1I, J). AIA activity was significantly impaired by blocking with O-Ag or IpaD, and increased slightly by blocking with IpaB (Figure 1K). Consistent with this, commercially available rabbit anti-IpaB polyclonal antibodies enhanced Shigella-dependent hemolysis of red blood cells *in vitro* (Supplemental Figure 2L). Preliminary experiments with IpaC did not reveal any impact on polyclonal serum function (data not shown); so this antigen was not investigated further. Taken together, these data indicate that at the polyclonal level, O-Ag specific antibodies contribute to SBA, OPA, and inhibition of adhesion and invasion, IpaD-specific antibodies inhibit adhesion and invasion, and IpaB-specific antibodies may enhance adhesion and invasion.

Although these studies revealed antigen-specific polyclonal antibody functions, it remained unclear how individual mAbs contribute to protection or potential disease enhancement. Such information is critical for vaccine design, as it can identify antigen epitopes and antibody germline sequences that drive protective immunity.^14–17^ We therefore undertook mAb discovery against O-Ag, IpaD, and IpaB.

### Antibodies against *S. flexneri* O-Ag are functionally diverse

To discover O-Ag-specific antibodies, cryopreserved PBMCs from four convalescent NHPs (Y01, Y02, 3a01, and 3a02) were stained with memory B cell markers (CD20^+^, IgG^+^) and OMVs from the infecting serotype (Y or 3a) labeled with the lipophilic dyes DiO and DiD. OMV double-positive memory B cells were sorted into 96-well PCR plates and paired native heavy and light chain sequences were recovered and cloned into Macaque IgG1 vectors for mAb production and validation (Figure 2A). Recombinant antibodies were screened for binding to intact bacteria of serotypes Y and 3a and negatively screened against a *S. flexneri* O-Ag knockout strain (Figure 2B). Across the four OMV sorts, we isolated 64 O-Ag-specific antibodies.

**Figure 2:**
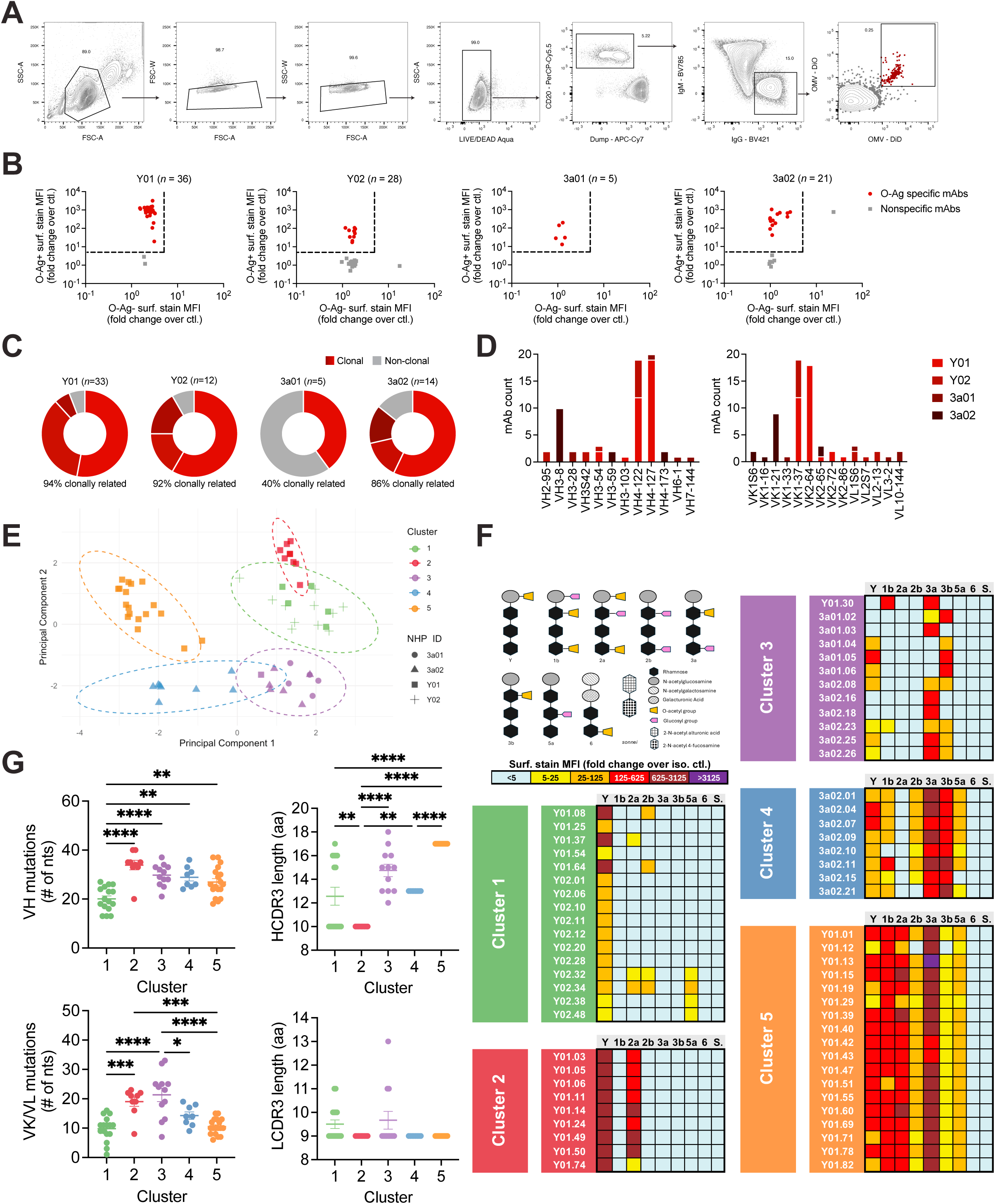
Anti-O-Ag monoclonal antibody discovery and cross-reactivity. (A) Gating strategy for the isolation of OMV-specific memory B cells from NHP PBMCs. (B) O-Ag specific mAbs as determined by fold change over isotype control of surface stain median fluorescence intensity (MFI) for Y/3a OAg^+^ and OAg^-^ strains. Number of mAbs assayed is shown above each graph. (C) O-Ag specific mAb clones (identical VH/VL pairings) for each NHP shown in shades of red; singletons in gray. Total mAbs per NHP indicated above each chart. (D) Frequencies of antibody V gene segments per NHP. (E) Principal component analysis (PCA) of log-transformed fold change over isotype control of surface stain MFIs against *S. flexneri* serotypes Y, 1b, 2a, 2b, 3a, 3b, 5a, & 6. mAbs binned by k-means clustering; ellipses show 95% CIs. (F) Schematic of O-Ag structures assayed, and heatmap of surface stain values for each mAb across all *S. flexneri* serotypes and *S. sonnei* (S.). Conducted in duplicate. (G) Mutation frequencies and CDR3 lengths of O-Ag specific mAbs by PCA cluster (n=8-18 mAbs per cluster). Ordinary one-way ANOVAs with Tukey’s multiple comparisons tests. \**p*< 0.05, \*\**p*< 0.01, \*\*\**p*< 0.001, and \*\*\*\**p*< 0.0001.

Within most NHPs, we observed a high proportion of clonally related sequences, consistent with prior reports that antibody responses against glycan targets are typically highly clonal (Figure 2C).^40,41^ Most germline gene pairings were unique to individual NHPs; however, VH4-122 and VK1-37 were shared between animals, comprising 36% of antibodies from Y01 and 58% from Y02 (Figure 2D). Altogether, the anti-O-Ag mAbs have on average 27 VH nucleotide substitutions (9.5%) and 14 VK/VL nucleotide substitutions (4.9%) (Supplemental Figure 3A), which is substantial for anti-polysaccharide antibodies.^42–44^ Furthermore, these mAbs have diverse CDR3 lengths: 10-18 amino acids for HCDR3s and 8-13 amino acids for LCDR3s (Supplemental Figure 3A). These features suggest that the antibodies were derived from a mature memory response shaped by repeated *Shigella* exposures. Together, these findings demonstrate that *S. flexneri* infection in NHPs elicits O-Ag-specific B cell responses characterized by significant somatic hypermutation, despite O-Ag being a T cell independent antigen.

O-Ag responses are generally considered to be serotype-specific, yet polyclonal sera sometimes exhibit cross-serotype reactivity.^45,46^ It is unclear whether this breadth arises from a mixture of strictly serotype-specific antibodies, or from the activity of cross-reactive O-Ag antibodies. To assess the serotype specificity of our O-Ag mAbs, we tested each antibody for binding to the surface of *S. flexneri* serotypes Y, 1b, 2a, 2b, 3a, 3b, 5a, 6, and *S. sonnei* (Figure 2F, Supplemental figure 3B). Principal component analysis of surface stain values grouped antibodies into five distinct functional clusters (Figure 2E, F). Cluster 1 comprised primarily Y-specific antibodies from NHPs Y01 and Y02 with limited cross-reactivity to 2a, 2b, and 5a. Cluster 2 consisted of Y- and 2a-cross reactive antibodies from Y01. Cluster 3 contained antibodies from Y01, 3a01, and 3a02, with primary specificity for 3a and/or 3b and minor reactivity to Y and 1b. Cluster 4 included relatively cross-reactive antibodies from 3a02 that bound Y, 1b, 2b, 3a, 3b, and 5a. Finally, cluster 5 contained the most broadly cross-reactive antibodies, isolated from Y01, which bound all tested *S. flexneri* serotypes except 6. Notably, none of the antibodies bound *S. flexneri* 6 nor *S. sonnei*, both of which have distinct O-Ag backbones (Figure 2F).

Sequence analysis revealed that cluster 1 antibodies, the least cross-reactive, displayed the lowest level of somatic hypermutation and some of the shortest CDR3s (Figure 2G). In contrast, antibodies from the more cross-reactive clusters tended to exhibit higher levels of somatic hypermutation and longer CDR3s, suggesting that broad cross-reactivity emerges only after extensive affinity maturation (Figure 2G). Taken together, these data demonstrate that anti-*Shigella* O-Ag antibodies span a spectrum of specificities. Antibodies in our library with a limited number of somatic mutations were more serotype-specific, while those exhibiting a greater degree of maturation could bind multiple *S. flexneri* serotypes, but not *S. flexneri* 6 or *S. sonnei*.

We next assessed effector functions of our O-Ag specific mAbs across our *S. flexneri* panel. Surface binding was tightly correlated with the ability to coordinate complement-mediated killing (Figure 3A, D, Supplemental Figure 4A). In contrast to SBA, antibody binding was not as strongly correlated with complement-independent opsonophagocytosis or T3SS inhibition (Figure 3B, C, E, F, Supplemental Figure 4B, C). While OPA and hemolysis assays resulted in statistically significant correlations with surface binding across most strains, they did not against serotype Y, and there were numerous exceptions in which mAbs bound the bacterial surface of a particular strain but did not have activity in OPA or hemolysis assays (Figure 3E, F, Supplemental Figure 4B, C). Furthermore, opsonophagocytic activity and T3SS inhibition were only weakly correlated with each other, and there were numerous instances in which mAbs coordinated one function but not the other (Figure 3G, Supplemental Figure 4D). Taken together, these results indicate that while surface binding of anti-O-Ag antibodies is generally sufficient to trigger complement-mediated bacteriolysis among *S. flexneri* strains, it is necessary but not sufficient to trigger complement-independent opsonophagocytosis and T3SS virulence inhibition, implying these functions are more sensitive to factors beyond an antibody’s ability to merely bind O-Ag, such as the distinct geometry or kinetics of antibody binding.

**Figure 3:**
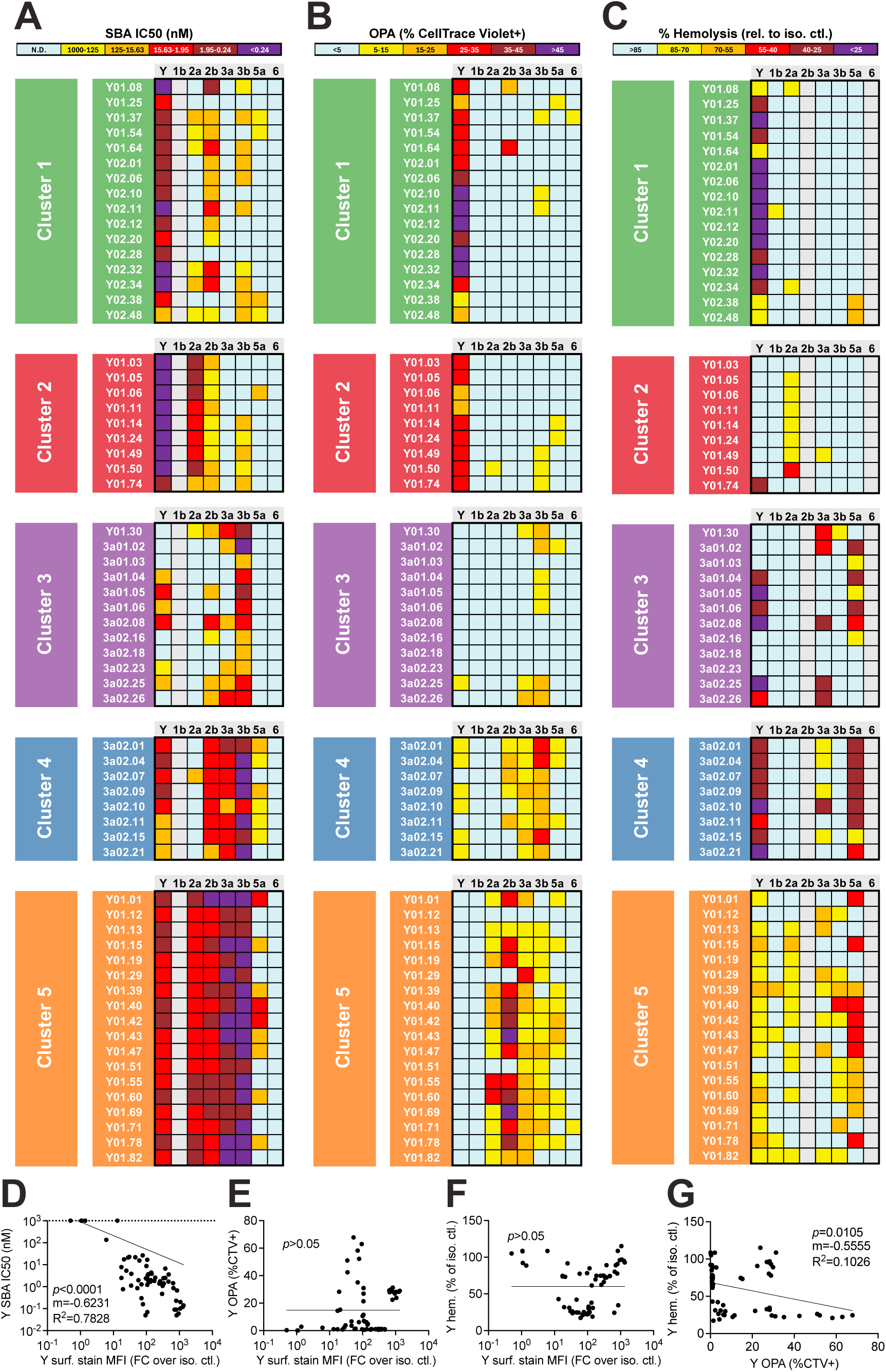
Functional activities of anti-O-Ag antibodies. (A) SBA IC_50_ values of each O-Ag specific mAb against serotypes Y, 2a, 2b, 3a, 3b, 5a, & 6. Our serotype 1b isolate was too serum sensitive to be used in SBA. Conducted in duplicate. (B) OPA activity of each mAb against serotype Y, 1b, 2a, 2b, 3a, 3b, 5a, & 6. Conducted in duplicate. (C) Percent hemolysis relative to isotype control (rel. to iso. ctl.) for each mAb against serotypes Y, 1b, 2a, 3a, 3b, & 5a. Our serotype 2b & 6 strains had no detectable hemolytic activity. Conducted in duplicate. (D-G) Correlations between serotype Y (D) surface stain and SBA IC_50_, (E) surface stain and OPA, (F) surface stain and hemolysis, and (G) OPA and hemolysis. Each data point represents a mAb. mAbs that had no detectable SBA activity were assigned an SBA IC_50_ of 1,000nM. For (D), log-log best fit line. For (E-G), semi-log best fit line. Extra sum-of-squares F Test to determine whether the slope significantly differs from 0.

### IpaD and IpaB elicit T cell responses in infected macaques

Antibodies are important components of acquired immunity against *Shigella* infection,^47^ but T cells also play an important role.^48^ As a polysaccharide, O-Ag cannot induce MHC-dependent T-cell activation; in contrast, the protein antigens IpaD and IpaB are efficiently processed and have been directly shown to elicit T cell responses in mice.^49^ Therefore, we investigated whether they elicited T cell responses in our NHP samples. PBMCs were stimulated *in vitro* with IpaD, IpaB, or the positive control superantigen *Staphylococcus* enterotoxin B (SEB),^50^ and activation was assayed by concentrations of T helper cytokines in media and by flow cytometry analysis of both surface activation markers and intracellular cytokines (Supplemental Figure 5A). Cytokine profiling of supernatants showed a weak, non-significant induction of IFN-γ and IL-17A, a strong induction of IL-21, IL-10, IL-6, and TNFα, and no effect on IL-2, IL-13, IL-4, or IL-5 (Supplemental Figure 5B). Flow cytometry revealed significant activation of CD4 and CD8 T cells by IpaD and IpaB, measured by the activation induced markers (AIM) OX40 and 4-1BB (Supplemental Figure 5C, D). Intracellular staining confirmed induction of IFN-γ in CD8 T cells, NK cells, and NKT cells, and induction of IL-17A in CD8 T cells and NK cells, but no significant induction in CD4 T cells. These results indicate that both IpaD and IpaB elicit cellular immune responses in NHPs, though with smaller effects on IFN-γ and IL-17A than expected based on previous reports.^48,49^

### Antibodies against the T3SS can both inhibit and enhance virulence

We next examined antibody responses against IpaD and IpaB. Following a workflow similar to the O-Ag-specific antibody discovery, cryopreserved PBMCs from six convalescent NHPs for IpaD (D02, D13c, D13r, D14, D16, and D17) and six for IpaB (B05, B07, B08, B10, B16, and B17) were stained with memory B cell markers (CD20^+^, IgG^+^). IpaD-specific B cells were identified using biotinylated IpaD complexed with AF488 and AF647-conjugated streptavidin as sorting baits, and IpaB-specific B cells were identified using IpaB labeled with AF488 and AF647-conjugated NHS-esters (Figure 4A). Antigen double-positive memory B cells were sorted into 96-well plates and paired heavy and light chain sequences were recovered (Figure 4A). Antibodies were screened by ELISA for antigen binding, and counter-screened against ovalbumin as a nonspecific control (Supplemental Figure 6A, B). Across the twelve Ipa protein sorts, we isolated 29 IpaD-specific antibodies and 24 IpaB-specific antibodies. Nearly all IpaD-specific antibodies were high affinity: 26 antibodies exhibited sub-nanomolar EC_50_ values by ELISA, while the remaining three had EC_50_s below 10 nM (Figure 4B). The IpaB antibodies exhibited a wide range of affinities – 11 were high affinity with sub-nanomolar ELISA EC_50_s, but the remainder had a wide range of affinities as low as 228 nM (Figure 4B). Both anti-IpaD and anti-IpaB antibodies were derived from diverse germline gene pairings. VH4-173 and VK1-37 were paired in both B08 and B17, but all other germline pairings were unique to each NHP (Figure 4C). Sequence analysis revealed comparable somatic hypermutation between the two libraries, averaging 24 VH nucleotide substitutions (8.1%) and 17 VK/VL nucleotides substitutions (6.0%) for IpaD-specific antibodies, and 21 VH nucleotide substitutions (7.3%) and 15 VK/VL nucleotide substitutions (5.3%) for IpaB-specific antibodies (Supplemental Figure 6C).

**Figure 4:**
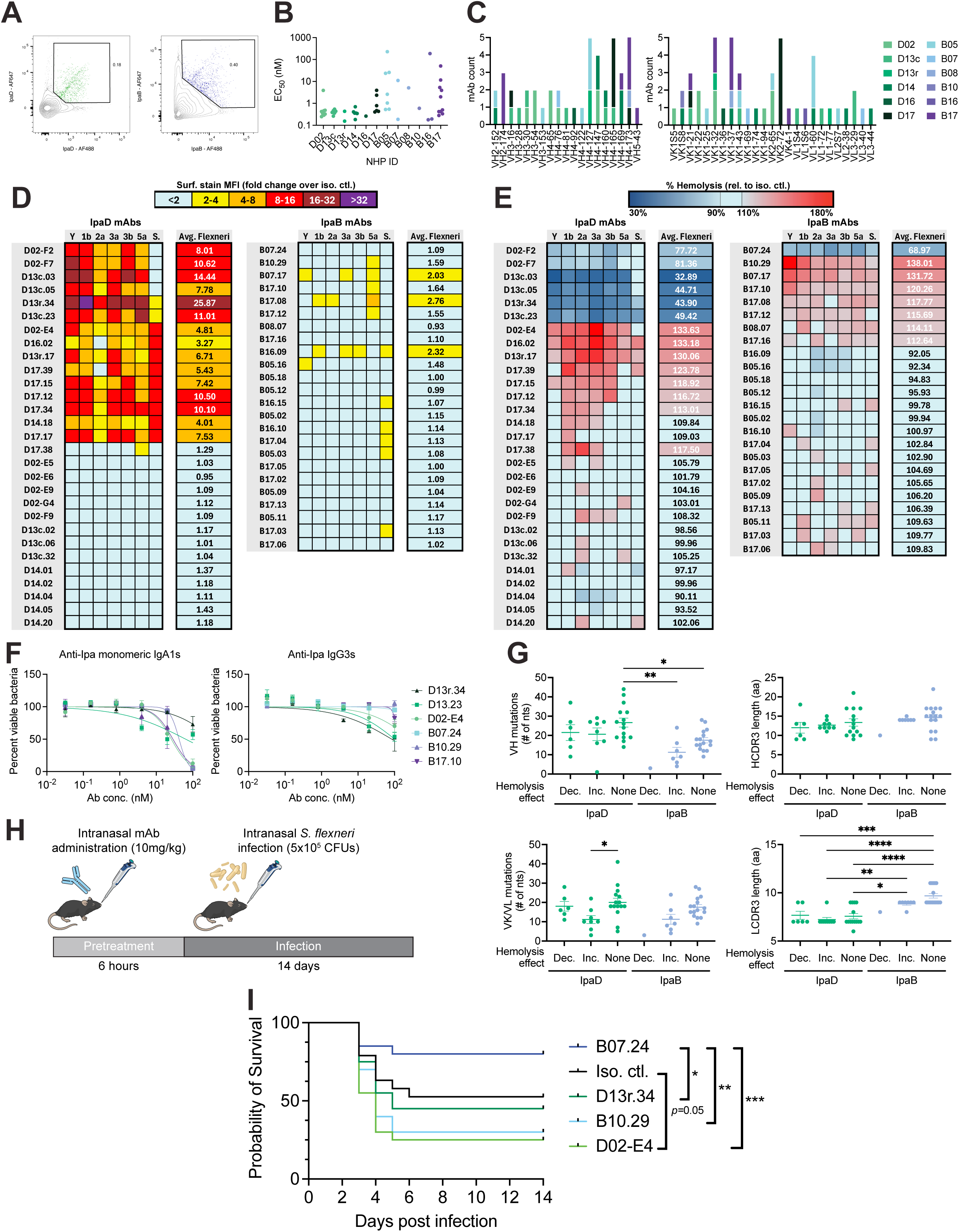
Anti-T3SS antibody discovery and functional activity. (A) Gating strategy for isolation of Ipa-specific memory B cells (CD20^+^, IgG^+^) from NHP PBMCs. (B) ELISA antigen EC_50_ values of Ipa-specific mAbs. (C) Frequencies of antibody V gene segments identified for each NHP. (D) Surface stain values for each Ipa-specific mAb against *S. flexneri* serotypes Y, 1b, 2a, 3a, 3b, & 5a, and against *S. sonnei* (S.). Bacteria were grown in the presence of 0.1% DOC for IpaB surface stains. (E) Percent hemolysis relative to isotype control of each Ipa-specific mAb against the same panel. (F) SBA curves of representative Ipa-specific mAbs in IgA1 monomer and IgG3 backbones. Conducted in duplicate three times, one representative experiment shown. (G) Mutation frequencies and CDR3 lengths of Ipa-specific mAbs, grouped by hemolysis effects (Dec.: decreases hemolysis, Inc.: increases hemolysis, None: no effect). (n=6-15 mAbs per group) Ordinary one-way ANOVAs with Tukey’s multiple comparisons tests. (H) Schematic of *in vivo* experiment. C57BL/6J mice were intranasally administered 10mg/kg of mAb, then infected 6 h later with 5 x 10^5^ CFU *S. flexneri* 2a (2457T). Mice were monitored daily for 14 days. (I) Survival of mice pretreated with each mAb. n = 20 per condition, two independent experiments combined. Log-rank analysis. \**p*< 0.05, \*\**p*< 0.01, \*\*\**p*< 0.001.

The T3SS needle tip consists of a conformationally dynamic complex including pentameric IpaD^51^ and (most likely) monomeric IpaB.^52^ Because all our mAbs were isolated using soluble, monomeric, recombinant protein, we assessed binding of these mAbs to native surface-exposed IpaD and IpaB on *S. flexneri* strains Y, 1b, 2a, 3a, 3b, and 5a (Supplemental Figure 6D). Fifteen antibodies bound to surface IpaD on at least five strains, with highly consistent binding patterns across serotypes (Figure 4D). In contrast, even with a modified surface stain protocol performed in the presence of deoxycholate (a bile salt known to recruit IpaB to the needle tip^52^) and amplified binding signal using a polyclonal tertiary antibody, IpaB specific antibodies exhibited weak and variable surface staining (Figure 4D). *S. sonnei* shares 97.3% IpaD and 99.0% IpaB sequence identities with *S. flexneri*, and both antigens have previously been shown to elicit cross-protective immune responses.^49^ Therefore, we also tested our antibody libraries for binding to *S. sonnei*. Eleven of the 15 *S. flexneri* IpaD binding antibodies also bound *S. sonnei*, while the IpaB antibody library exhibited similarly weak and inconsistent binding *to S. sonnei* (Figure 4D).

Functional assays revealed that IpaD-specific antibodies lacking surface binding had no impact on T3SS function (Figure 4E). Nearly all surface-binding IpaD antibodies impacted hemolysis, but surprisingly much of the impact appeared harmful: six blocked hemolysis, but eight enhanced it (Figure 4E). Notably, four of the blockers and five of the enhancers had no effect on *S. sonnei*, demonstrating that cross-species functional activity is limited at the monoclonal level (Figure 4E). Despite the inconsistent IpaB surface staining, hemolysis assays revealed consistent effects across both *S. flexneri* and *S. sonnei*: one IpaB-specific antibody reduced hemolysis, seven enhanced it, and the remainder had no effect (Figure 4E).

Both anti-IpaD and anti-IpaB IgG1 antibodies did not mediate SBA nor OPA against *S. flexneri* serotype Y (Supplemental Figure 6E, F). Given that these functions are isotype-dependent, we re-cloned a subset of representative antibodies into IgG3 and monomeric IgA1 backbones – isotypes shown to have more potent functions in similar assays compared to IgG1.^53,54^ Isotype switched antibodies did not exhibit OPA activity, but some did weakly coordinate complement-mediated killing (Figure 4F, Supplemental Figure 6G).

Sequence analysis revealed no major differences in somatic hypermutation nor CDR3 lengths between antibodies with distinct functional profiles within each antigen-specific library, suggesting that antibody-mediated functional modulation of T3SS activity does not require unusually complex maturation pathways (Figure 4G). Notably, IpaD antibodies tended to have more VH mutations while IpaB antibodies tended to have longer LCDR3s (Figure 4G). Collectively, these findings indicate that IpaD and IpaB responses in NHPs are polyclonal and diverse but consistently affinity-matured, and they can modulate T3SS activity across serotypes and can weakly coordinate complement-mediated killing as IgG3 and monomeric IgA1 antibodies.

To our knowledge, only one anti-*Shigella* T3SS mAb, the anti-IpaD VHH IpaD-318, has been evaluated *in vivo*, using a murine intranasal challenge model.^55^ IpaD-318 enhanced hemolysis *in vitro*, but paradoxically protected mice from *Shigella* challenge *in vivo*. Given this context, we next tested whether our natural infection-derived antibodies were sufficient to impact *in vivo* disease, and whether the direction of that impact was consistent with our *in vitro* results. Five antibodies – two hemolysis blockers (D13r.34 & B07.24), two hemolysis enhancers (D02-E4 & B10.29), and one isotype control (VRC01, an antibody against human immunodeficiency virus GP120^56^) – were administered intranasally (10 mg/kg) to C57BL/6J mice 6 h prior to LD50 challenge (5 x 10^5^ CFUs *S. flexneri* 2a) (Figure 4O). Among the *in vitro* blockers D13r.34 and B07.24, D13r.34 had no effect, while B07.24 significantly improved survival compared to D13r.34, B10.29, and D02-E4. Among the *in vitro* enhancer mAbs D02-E4 and B10.29, B10.29 trended toward decreased survival, and D02-E4 significantly decreased survival compared to B07.24 and the isotype control (Figure 4P). Thus, anti-IpaD and anti-IpaB mAbs are sufficient to modulate disease in a murine intranasal challenge model, and the direction of that modulation is generally consistent between *in vitro* and *in vivo* assays.

### Anti-IpaD and anti-IpaB antibody functions map to specific epitopes

Antibodies against a given target may have opposite functional effects either because they bind a different epitope, or because they trigger different conformational changes when binding to the same epitope. Therefore, given the opposite functional effects observed for the mAbs against both IpaD and IpaB, we next examined whether these functional differences may reflect differences in epitope specificity. Competition ELISAs revealed that functional IpaD antibodies clustered into two hemolysis-blocking epitopes (epitopes D-III and D-Vb) and one enhancing epitope (epitope D-IVb) (Figure 5A). Most IpaB hemolysis-enhancing antibodies clustered into one dominant epitope (epitope B-I), while the most potent mAbs B10.29 (the strongest enhancer) and B07.24 (the only blocker) both spanned two otherwise non-functional epitopes (epitopes B-II and B-IIIa) (Figure 5B).

**Figure 5:**
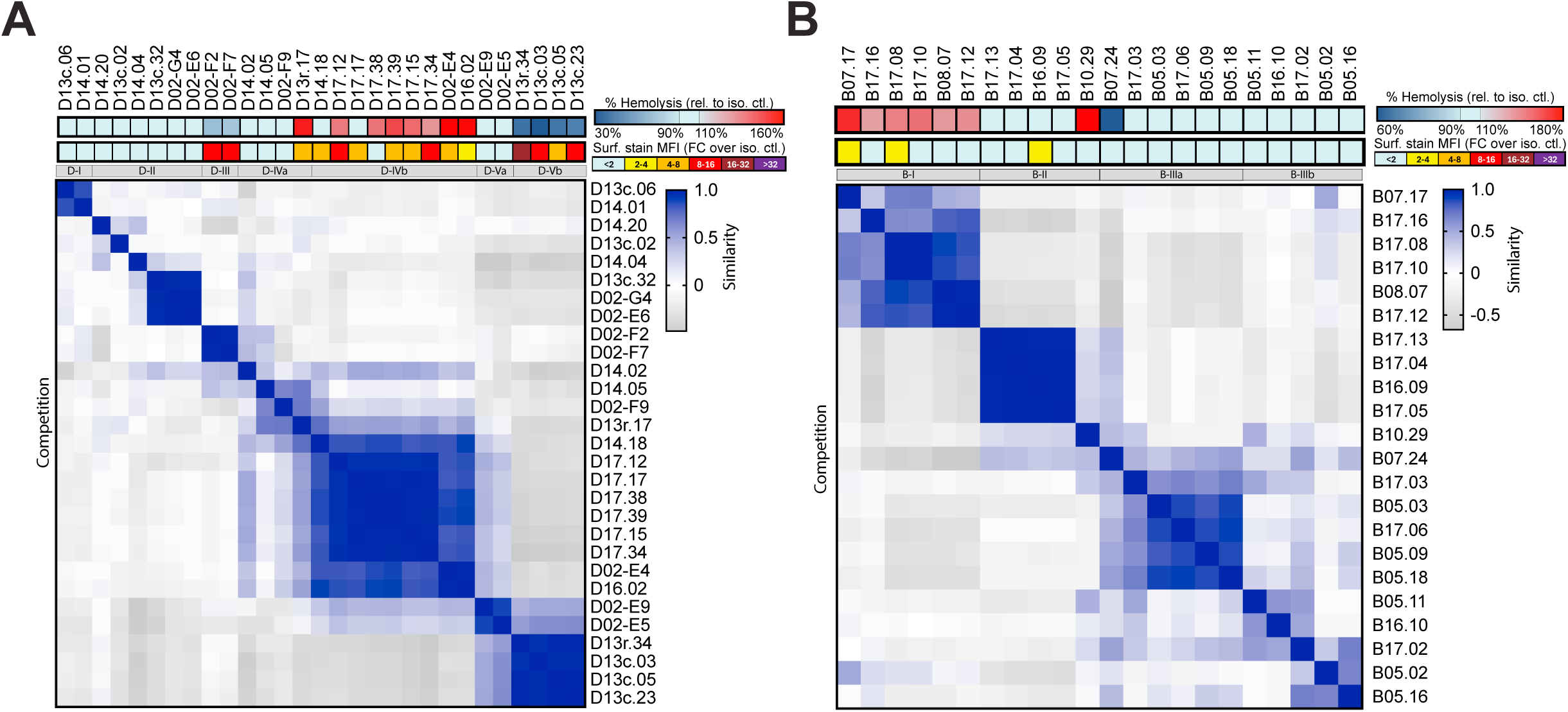
Epitope binning of IpaD and IpaB antibodies. (A-B) Competition ELISA of (A) IpaD-specific mAbs and (B) IpaB-specific mAbs. Values normalized to unblocked and self-blocking controls. Conducted in duplicate. Pearson-correlation similarity matrix applied. Average surface stain and hemolysis values of each mAb relative to isotype control across all *flexneri* serotypes assayed is indicated above.

To identify these functional epitopes on both IpaD and IpaB, we sought to determine representative mAb:antigen complex structures by cryogenic electron microscopy (cryo-EM). The structure of full-length IpaB has not yet been solved and our attempts at IpaB structural determination were unsuccessful. However, we did determine that the strongest enhancer antibody, B10.29, was the only antibody capable of binding in an ELISA to IpaB_28.226_, a domain essential for IpaB-IpaD binding^57^ (Supplemental Figure 7A). IpaD has a well-defined structure with eight a-helices and five b-strands arranged in three domains: the N-terminal domain, the central coiled-coil domain, and the distal domain^58^ (Supplementary Figure 7B and 7C). Complexes between IpaD and two representative hemolysis blocking mAbs (D02-F2 and D13r.34) and one hemolysis enhancing mAb (D02-E4) were resolved in two different structures: one with IpaD in complex with D13r.34, and another with IpaD in complex with both D02-F2 and D02-E4 (Table 1, Figure 6A, 6B, Supplemental Figures 8 & 9).

**Figure 6:**
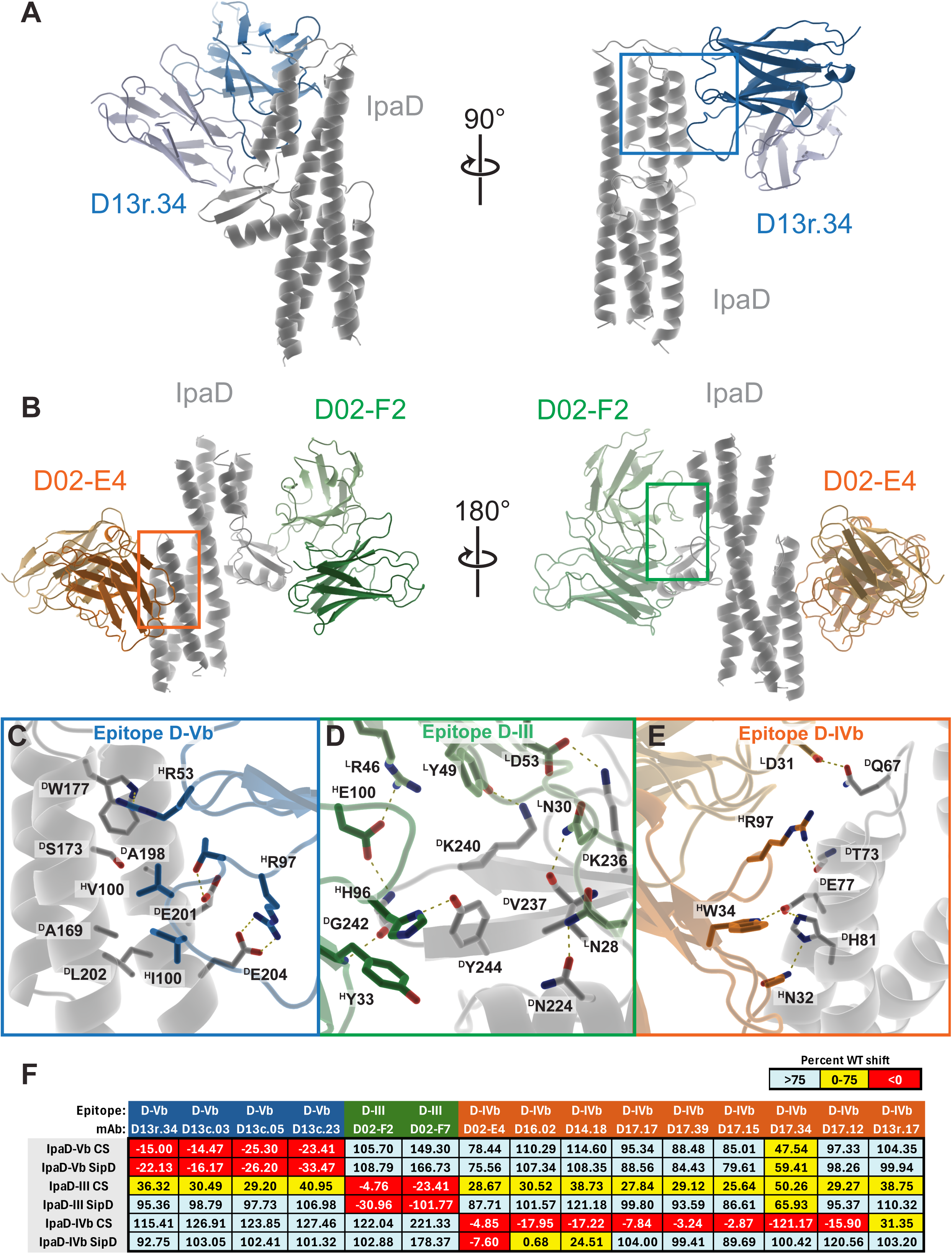
Determination of IpaD-specific antibody binding epitopes. (A) Cryo-EM structure of IpaD (gray) in complex with D13r.34 (blue). The mAb light chain is depicted in a lighter color. The blue box shows the region depicted more closely in panel C. (B) Cryo-EM structure of IpaD (gray) in complex with D02-F2 (green) and D02-E4 (orange). The mAb light chains are depicted in a lighter color, and the green and orange boxes show regions depicted more closely in panels D and E, respectively. (C-E) Depictions of the Fab-IpaD interface for D13r.34 (blue) (C), D02-F2 (green) (D), and D02-E4 (orange) (E), showing key interactions. Residues are prefaced with a “D” if they are part of IpaD, “H” if they belong to the mAb heavy chain, and “L” if they belong to the mAb light chain. Likely electrostatic interactions are shown as yellow dashed lines. Light chains are depicted in a lighter shade than heavy chains, and IpaD is gray. (F) Biolayer interferometry analysis showing relative binding of each antibody to mutagenized IpaD constructs. Data are expressed as percent binding signal (shift) normalized to WT IpaD shift for each antibody.

**Table 1:**
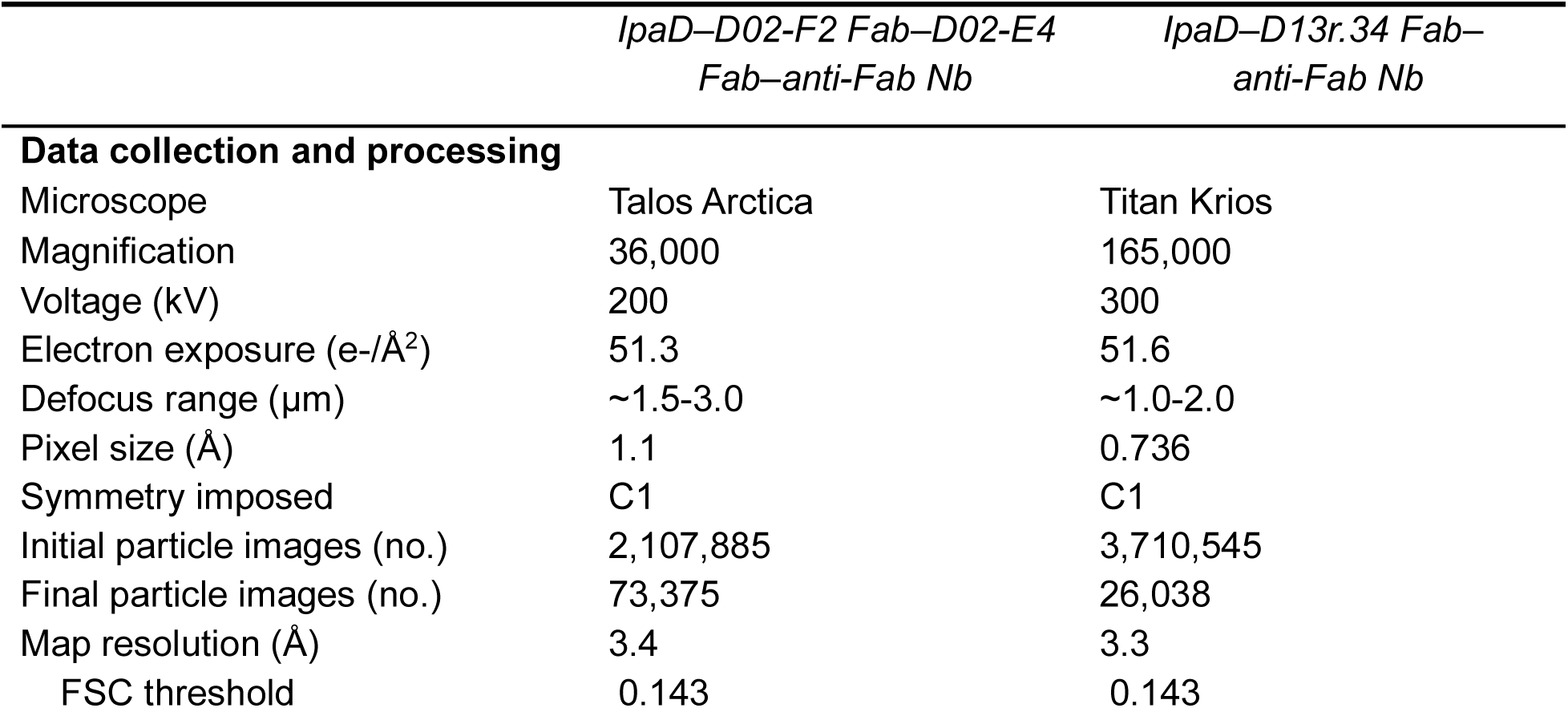
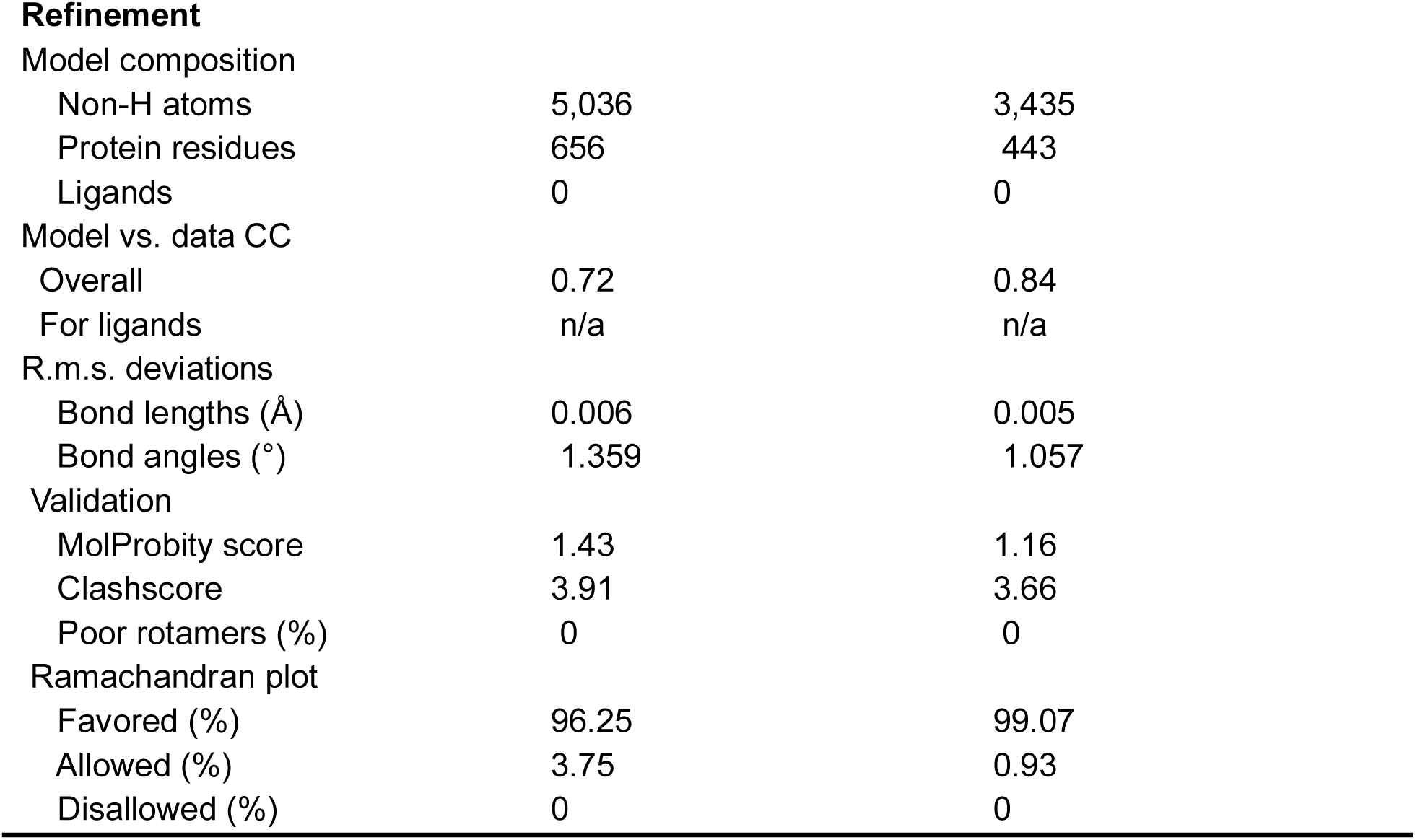
<u>Cryo-EM</u> data collection, refinement, and validation statistics.

|  | *IpaD–D02-F2 Fab–D02-E4*<br>*Fab–anti-Fab Nb* | *IpaD–D13r.34 Fab–*<br>*anti-Fab Nb* |
| --- | --- | --- |
| **Data collection and processing** |  |  |
| Microscope | Talos Arctica | Titan Krios |
| Magnification | 36,000 | 165,000 |
| Voltage (kV) | 200 | 300 |
| Electron exposure (e-/Å<sup>2</sup>) | 51.3 | 51.6 |
| Defocus range (μm) | ~1.5-3.0 | ~1.0-2.0 |
| Pixel size (Å) | 1.1 | 0.736 |
| Symmetry imposed | C1 | C1 |
| Initial particle images (no.) | 2,107,885 | 3,710,545 |
| Final particle images (no.) | 73,375 | 26,038 |
| Map resolution (Å) | 3.4 | 3.3 |
| FSC threshold | 0.143 | 0.143 |
| Refinement |  |  |
| Model composition |  |  |
| Non-H atoms | 5,036 | 3,435 |
| Protein residues | 656 | 443 |
| Ligands | 0 | 0 |
| Model vs. data CC |  |  |
| Overall | 0.72 | 0.84 |
| For ligands | n/a | n/a |
| R.m.s. deviations |  |  |
| Bond lengths (Å) | 0.006 | 0.005 |
| Bond angles (°) | 1.359 | 1.057 |
| Validation |  |  |
| MolProbity score | 1.43 | 1.16 |
| Clashscore | 3.91 | 3.66 |
| Poor rotamers (%) | 0 | 0 |
| Ramachandran plot |  |  |
| Favored (%) | 96.25 | 99.07 |
| Allowed (%) | 3.75 | 0.93 |
| Disallowed (%) | 0 | 0 |

Our structures revealed unambiguous identification of mAb binding sites at the domain level. D13r.34 binds a3 of the central coiled-coil domain and a4 of the distal domain in a manner that appears to be primarily dependent on its CDRH3 (Figure 6A, 6C). This epitope D-Vb is nearly identical to those previously reported for JPS-G3 and 20-IpaD, camelid VHH domains that inhibit *Shigella* hemolysis.^59^ In contrast, D02-F2 and D02-E4 bind regions of IpaD not recognized by previously reported IpaD mAbs.^55,59^ Both the heavy and light chains of D02-F2 are in close contact with epitope D-III, the distal domain at bC, bD, a5, and bE (Figure 6B, 6D). In contrast, the hemolysis enhancing mAb D02-E4 associated closely with epitope D-IVb, the N-terminal domain at a1 and a2, again with both heavy and light chains likely playing key roles in IpaD recognition (Figure 6B, 6E).

To validate these epitopes and lay a foundation for rational antigen design, we introduced targeted mutations within the D-Vb, D-III, and D-IVb epitopes. The mutations were either conservative substitutions with homologous *Salmonella* SipD residues, or more disruptive charge-swap (CS) mutations (Supplemental Figure 10A). Biolayer interferometry confirmed that both IpaD-Vb variants (SipD and CS) abolished epitope-specific binding (Figure 6F, Supplemental Figure 10C, D). The IpaD-III SipD variant abolished epitope-specific binding, while the IpaD-III CS variant disrupted all antibody binding (Figure 6E, Supplemental Figure 10E, F). The IpaD-IVb SipD variant only ablated D02-E4 binding, whereas the IpaD-IVb CS variant also blocked binding of other antibodies targeting that site (Figure 6E, Supplemental Figure 10G, H). Altogether, these data demonstrate that functional differences among anti-Ipa protein mAbs are consequences of distinct binding epitopes, and our preliminary epitope-specific IpaD mutagenesis suggests that rational design of IpaD variants may enable selective ablation of disease-enhancing antibody responses while preserving those that lead to inhibition of hemolysis.

## Discussion

*Shigella* infection is the second most common cause of diarrhea globally,^11^ and as rates of antimicrobial resistance continue to rise,^3–6^ an effective vaccine is increasingly important. Previous work in NHP challenge models^18^ and humans^21,22^ suggested that acquired immunity to *Shigella* infection is limited in breadth (often being serotype-specific) and duration. More recently, systems-level analysis of clinical samples supports roles for candidate vaccine antigens such as O-Ag^23,24,60^ and/or T3SS proteins^26–28^ in protection. While such data are critical and have informed the design of the current vaccine pipeline,^7^ these studies necessarily provide only correlative associations between immunological metrics and disease – they do not provide sufficient resolution or mechanistic insight to comprehensively inform the design of vaccine antigens. Here, we leveraged a unique opportunity provided by a natural outbreak of circulating *S. flexneri* strains in an NHP research colony to dissect antibody responses against candidate vaccine antigens (O-Ag, IpaB, and IpaD), with the aim of better understanding how to design effective *Shigella* vaccines. This outbreak provided novel insights into the primate immune response following repeated, low-dose exposures to contemporary circulating strains like those experienced in endemic contexts.

The key finding from this anti-O-Ag mAb discovery campaign was that a subset of these mAbs have remarkable breadth, with the ability to bind and coordinate complement-mediated bacteriolysis against many *S. flexneri* serotypes (Figure 2F and 3A). While O-Ag is the most common antigen in candidate *Shigella* vaccines due to strong evidence for its importance in infection^7,18–22^, its high structural variability between serotypes^61^ makes developing an affordable O-Ag vaccine with clinically relevant breadth challenging.^7^ Recent work has shown that immunization with OMVs from as few as four *Shigella* strains can provide broad, O-Ag dependent serum bactericidal responses in rats and rabbits, though not humans.^62,63^ An open question is whether broad and potent O-Ag responses consist primarily of many unique antibody lineages specific for a narrow range of serotypes, or a few lineages that are cross-reactive. Here we find clonally related, broadly reactive mAbs that have undergone extensive (for a T cell independent antigen such as O-Ag) affinity maturation (Figure 2G). These data predict it is possible to produce such mAbs following repeated *Shigella* infections, and that vaccination strategies based on serial administration of specific serotypes may be more successful than a single multivalent formulation. However, the low clonal diversity and relatively high number of variable domain mutations in broad mAbs (Figure 2G) suggest that such antibodies may be challenging to elicit. Indeed, we are aware of only one other study reporting such levels of somatic hypermutation in anti-O-Ag mAbs^64^ (against *Klebsiella pneumoniae*). A recent analysis of mAbs isolated from humans immunized with a candidate *S. sonnei* vaccine showed levels of somatic hypermutation like those of the serotype-specific cluster 1 mAbs reported here,^65^ suggesting that alternative vaccine dose schedules or formulations could be required to elicit broad antibodies against O-Ag. This is of particular importance when considering that anti-polysaccharide antibodies are hardest to elicit in young children and elderly people,^66,67^ which are disproportionately affected by *Shigella* infection.^11^

While most IpaD- and IpaB-specific antibodies we isolated were non-functional, both antigens could elicit mAbs that potently inhibit hemolysis with relatively few mutations in a diverse set of germline genes (Figure 4C and 4G) and induce cellular immunity characterized by at least moderate IFNγ and IL-17A expression (Supplemental Figure 5). In contrast to the variability of O-Ag-specific mAbs, T3SS-directed antibodies were more uniformly cross-reactive against *S. flexneri* serotypes, and several exhibited cross-species activity against *S. sonnei.* However, we also observed evidence of *in vivo* antibody-dependent enhancement (ADE) of disease^68^ (Figure 4I) from mAbs against IpaD and IpaB in a murine challenge model. ADE has been most notoriously associated with viral pathogens such as Dengue virus^69,70^ but has also been reported for bacterial pathogens such as *Acinetobacter baumannii,*^71^ *Neisseria gonorrhoeae,*^72–74^ and *Pseudomonas aeruginosa.*^75^ While there have been reports of mAbs against IpaC^76^ or IpaD^55^ that enhance virulence *in vitro*, to our knowledge this is the first example of mAbs capable of increasing *Shigella* virulence *in vivo*. These data suggest that ADE could be another immune evasion mechanism for *Shigella*, which is already known to employ strategies that suppress innate^77,78^ and T cell responses.^79,80^ Importantly, protective and disease-enhancing effects mapped onto distinct epitopes of both proteins (Figure 5A and 5B). The strongest enhancing IpaB-specific mAb (B10.29) bound to the IpaD-binding N-terminal domain, suggesting it may function by modulating IpaD-IpaB interactions.^57^ Cryo-EM analysis of IpaD-mAb complexes revealed that protective mAbs primarily bind the distal domain (the least conserved part of the protein, suggestive of selective pressure), while the disease enhancing mAbs bind to the N-terminal domain (Figure 6A-E). Importantly, the distal domain of IpaD moves as the T3SS complex is primed for invasion,^51,81^ and a structure of the T3SS needle tip complex from *Salmonella* shows a SipD (the *Salmonella* homolog to IpaD) pentamer in a conformation that would not accommodate the N-terminus in the position observed in monomeric crystal structures.^82^ Thus, it is tempting to speculate that within the assembled T3SS needle tip, IpaD’s N-terminal and distal domains sample conformational states that have different capacities for host cell invasion, analogous to what is observed for pre-fusion and post-fusion conformations in the fusion proteins of many viral pathogens.^14,83–86^ If future work determines this is true, it will suggest that the different functional effects of mAbs against these domains may be a result of their ability to stabilize different IpaD conformational states. Moreover, by mutating IpaD we specifically ablated antibody binding to each epitope (Figure 6F), suggesting that IpaD can be rationally designed as a vaccine immunogen that does not elicit antibodies targeting disease enhancing epitopes. Taken together, these data indicate that Ipa proteins hold promise as vaccine antigens and can be rationally designed to prevent ADE of virulence.

Altogether, this extensive mAb discovery campaign is an important first step towards equipping vaccinologists with immunological and structural data that can support the rational design of *Shigella* vaccine candidates. Future work should focus on additional structural insights into anti-*Shigella* mAb function, and determine if the extensive somatic hypermutation for broad anti-O-Ag mAbs and disease enhancing mAbs against T3SS antigens observed here in NHPs can occur in humans. Our findings suggest that integrating structural and immunological data into *Shigella* vaccine design strategies can likely overcome existing challenges to create broad and efficacious vaccines that incorporate multiple antigens.

## Acknowledgements

We thank Finora Franck for invaluable program management support over the course of this project. This work was supported by the Wellcome Trust RESISTANT grant, and we thank past and present members of the RESISTANT Research Steering Group (including Wellcome Trust Program Officers) for their invaluable support of this work, including Dr. Pete Gardner, Dr. Colleen Loynachan, Dr. Kate Mills, Dr. Nathalie Herze-Vourc’h, and Dr. Christiane Gerke. We also thank Dr. Marcela Pasetti for supplying the recombinant IpaC and IpaB used in initial serological assays. We also thank IAVI colleagues Dr. Elise Landais, Dr. Jon Heinrichs, and Dr. Tawanda Mandizvo for a critical reading of this manuscript. Cryo-EM data were collected at the Harvard Cryo-EM Center for Structural Biology at Harvard Medical School. Structural biology applications used in this project were compiled and configured by SBGrid. M.S.A.G. is supported by the HHMI Hanna H. Gray Fellows Program. We thank members of the Harvard Cryo-EM Center for Structural Biology and SBGrid for their support. We would like to thank Cheryl Kim, Denise Hinz, and the team at the La Jolla Institute for Immunology Flow Cytometry Core Facility for their assistance with data collection. We thank the WNPRC Colony Services, Veterinary Services, and SPI Units for their care of the NHPs and sample collection. Additionally, we thank Dr. Heather Simmons for her support and advice, and Dr. Simmons, Dr. Puja Basu, Nicole Oldenburger, Abby Bradford, Alanna Friscino, Jessie Kelly, and Chris Huffman for their sample processing, necropsy, and isolate shipment efforts. We also thank the University of Wisconsin-Madison Veterinary Medical Teaching Hospital Clinical Pathology Laboratory for their culture and susceptibility performance. Research reported in this publication was supported in part by the Office of the Director, National Institutes of Health under Award Number P51OD011106 to the Wisconsin National Primate Research Center, University of Wisconsin-Madison. This research was conducted in part at a facility constructed with support from Research Facilities Improvement Program grant numbers RR15459-01 and RR020141-01. The content is solely the responsibility of the authors and does not necessarily represent the official views of the National Institutes of Health.

## Data Availability

The atomic coordinates for both structures will be deposited at the Protein Data Bank and corresponding cryo-EM maps will be deposited at the Electron Microscopy Data. Genome and antibody sequences will be made available on NCBI.

---

**Supplemental Figure 1:**
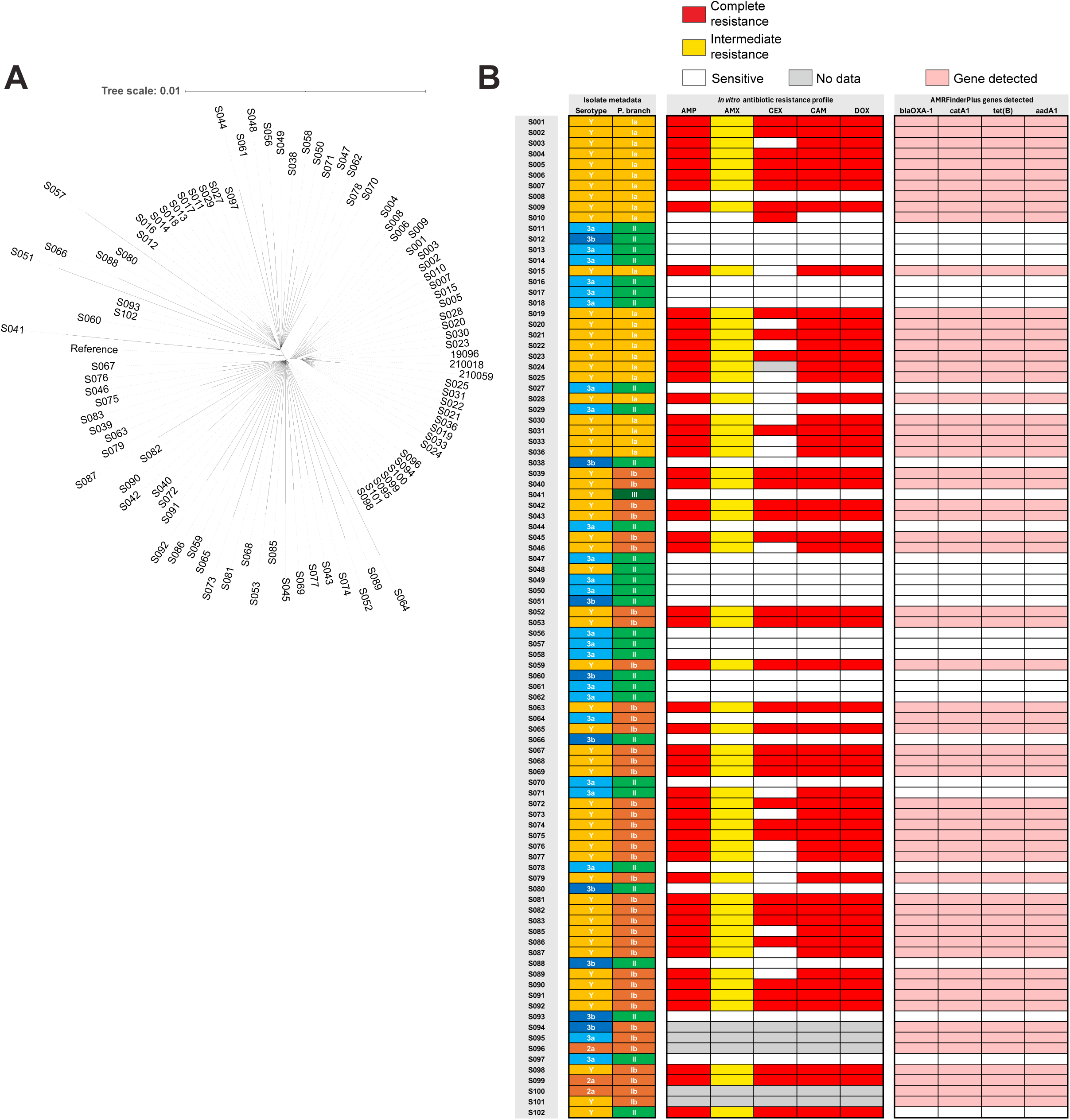
Bacterial phylogeny and antimicrobial resistance, related to Figure 1. (A) Unrooted phylogenetic tree of the same strains shown in Figure 1B. Branch lengths are proportional to evolutionary distance. (B) Heatmap of serotype, phylogenetic branch, antibiotic resistance profiles, and detected antibiotic resistance genes for each strain. AMP = ampicillin (MIC ≥ 32 µg/ml = resistant); AMX = amoxicillin (MIC = 16 µg/ml = intermediate); CEX = cephalexin (MIC ≥ 8 µg/ml = resistant); CAM = chloramphenicol (MIC ≥ 64 µg/ml = resistant); DOX = doxycycline (MIC = 16 µg/ml = resistant).

**Supplemental Figure 2:**
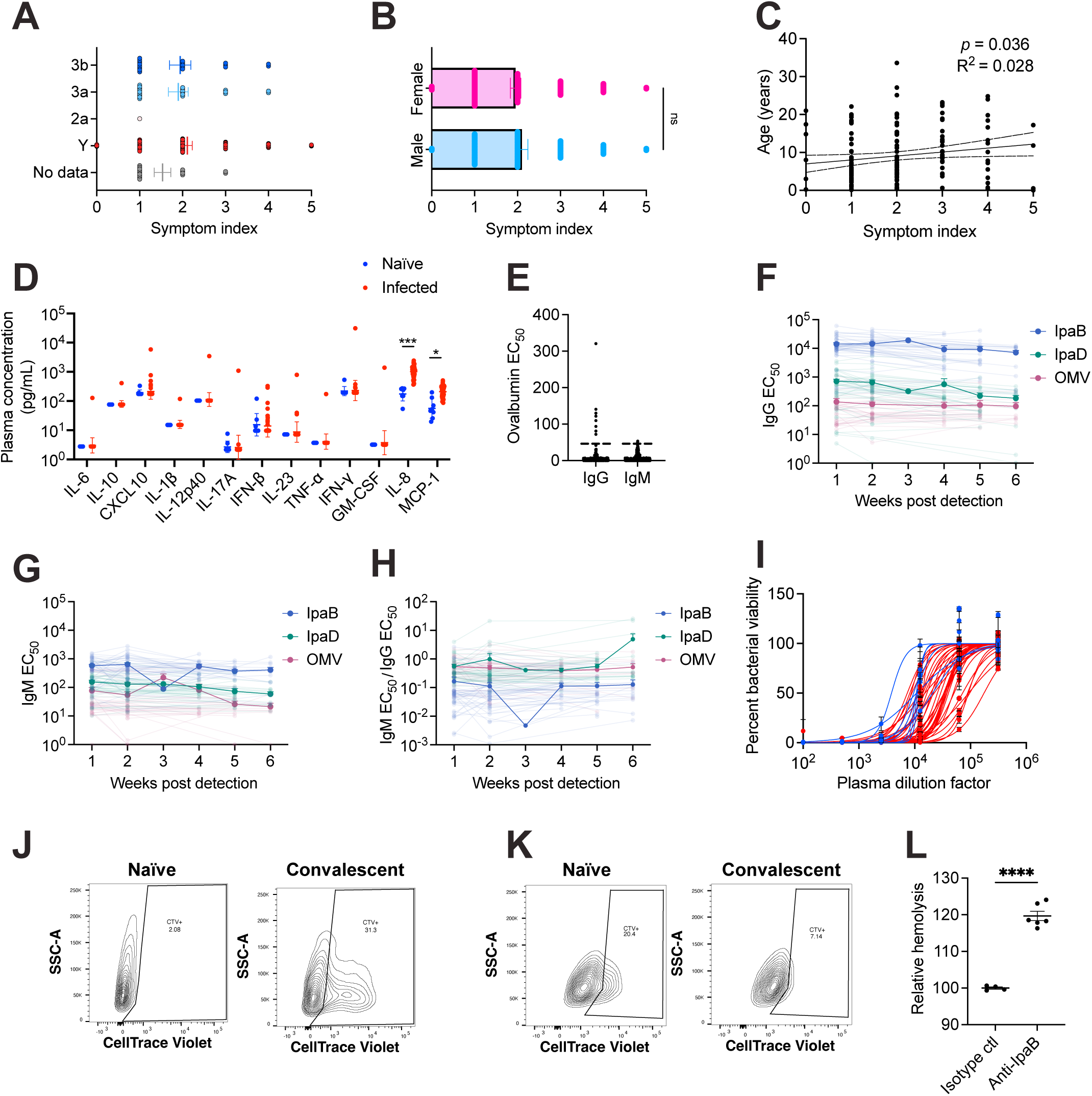
NHP clinical outcomes and serology, related to Figure 1. (A) Symptom index by infecting serotype (n=1-109). One-way ANOVA with Tukey’s multiple comparisons test. Mean ± SEM. (B) Symptom index by sex (n=68 males, n=91 females). Unpaired t-test with Welch’s correction. (C) Correlation of symptom index and age (n=158). Simple linear regression with 95% CI. (D) Cytokine levels of naïve (n=8) and 1-week post detection of infection plasma (n=41). Geometric mean ± geometric SD, multiple lognormal Welch’s t-tests with Holm-Šídák correction. Conducted in duplicate. (E) IgG and IgM ELISA EC_50_ values against ovalbumin; samples > 40 excluded from ELISA titer analyses as polyreactive. (F-G) Time-course of serum (E) IgG and (F) IgM ELISA EC_50_ values from Y-infected NHPs against IpaB, IpaD, and Y OMVs (n=46). (H) Ratio of IgM / IgG EC_50_ values from (E-F) (n=46). (I) SBA curves corresponding to the serum bactericidal assay IC_50_ values plotted in Figure 1E. (J) Representative flow plots of opsonophagocytosis assays using naïve and convalescent plasma. THP-1 cells are identified by size, then cells that have internalized bacteria are identified by CellTrace Violet signal (CTV+). (K) Representative flow plots of adhesion and invasion assays using naïve and convalescent plasma. HeLa cells are identified by size, then the extent to which bacteria have adhered/invaded is measured by % of cells that are CellTrace Violet positive. (L) Percent hemolysis relative to isotype control of polyclonal IpaB antibodies against *S. flexneri* 2a (n=4-6 per condition, two independent experiments combined). Unpaired t-test with Welch’s correction. \**p*< 0.05, \*\*\**p*< 0.001, and \*\*\*\**p*< 0.0001.

**Supplemental Figure 3:**
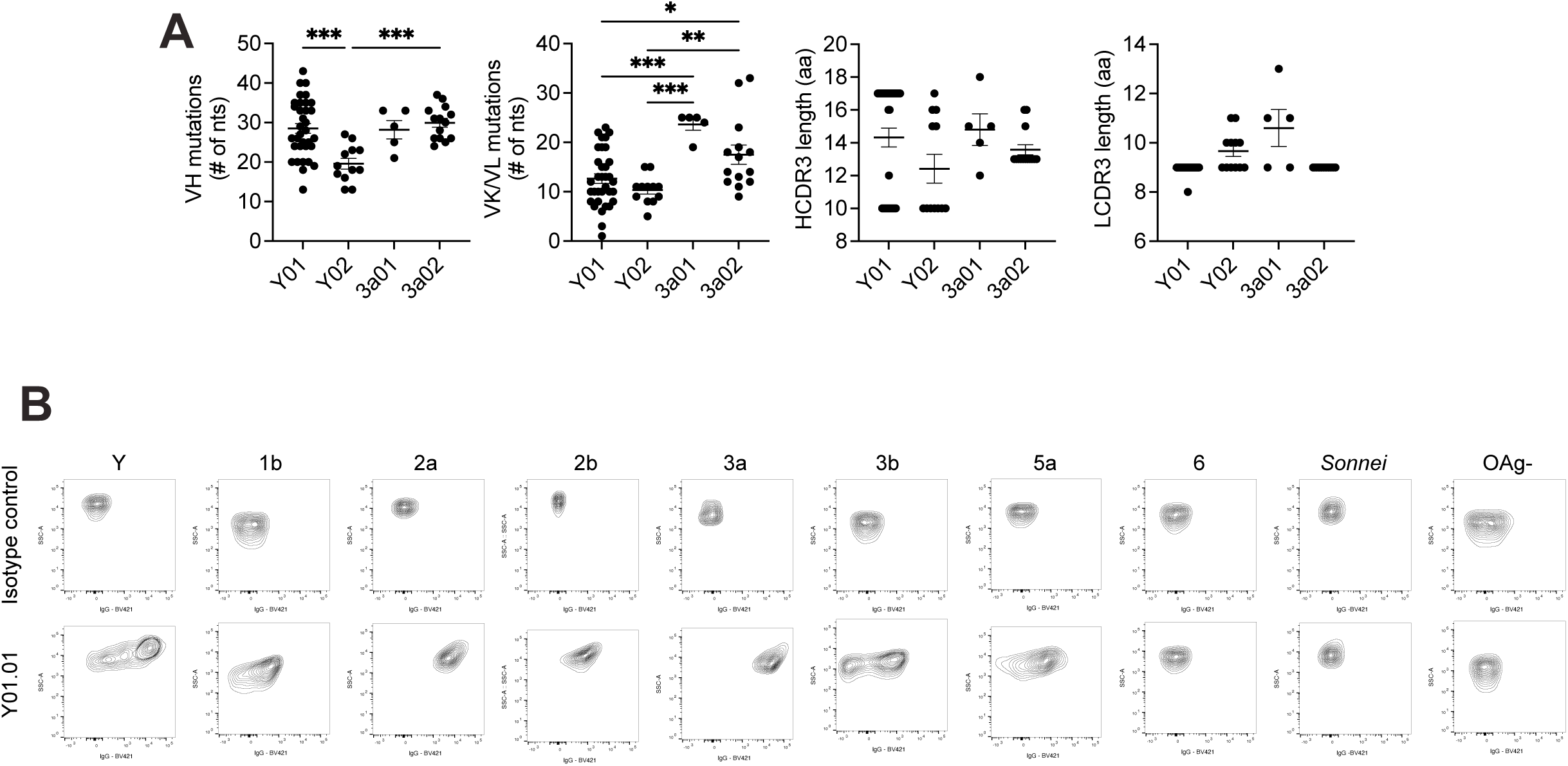
O-Ag mAb characterization, related to Figure 2. (A) Same data as Figure 2G. Mutation frequencies and CDR3 lengths of OAg-specific mAbs by NHP. (n=5-34 mAbs per NHP) Ordinary one-way ANOVAs with Tukey’s post-test. (B) Representative flow plots of bacterial surface stains against *S. flexneri* Y, 1b, 2a, 2b, 3a, 3b, 5a, 6, OAg^-^, and *S. sonnei*. \**p*< 0.05, \*\**p<0.01*, \*\*\**p*< 0.001, and \*\*\*\**p*< 0.0001.

**Supplemental Figure 4:**
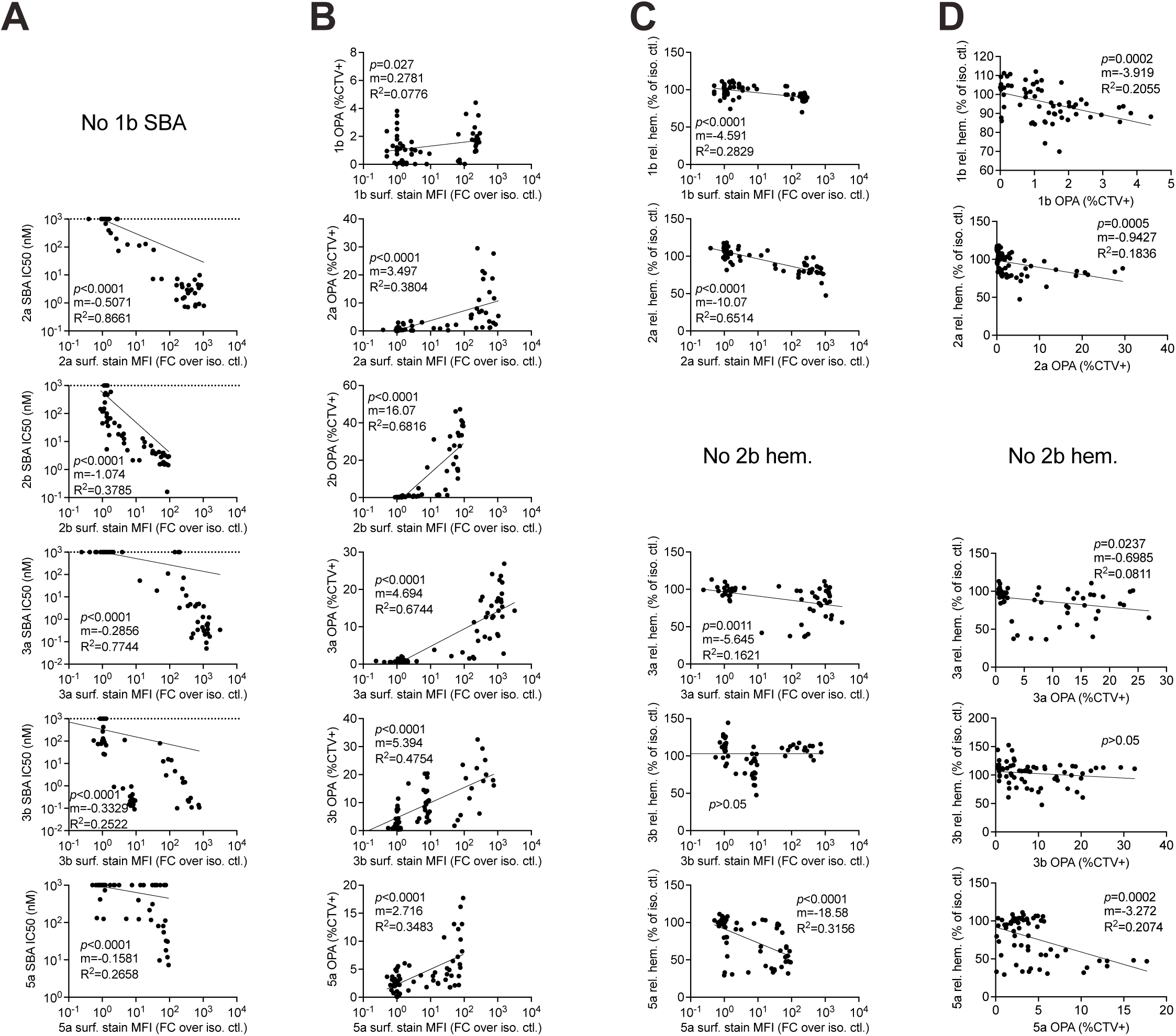
Functional correlations of O-Ag mAbs, related to Figure 3. (A-D) Correlations per serotype between (A) surface stain and SBA IC_50_, (B) surface stain and OPA, (C) surface stain and hemolysis, and (D) OPA and hemolysis. Each data point represents a mAb. mAbs that had no detectable SBA activity were assigned an SBA IC_50_ of 1000nM. Log-or semi-log best fit lines, Extra sum-of-squares F Test to determine whether the slope significantly differs from 0.

**Supplemental Figure 5:**
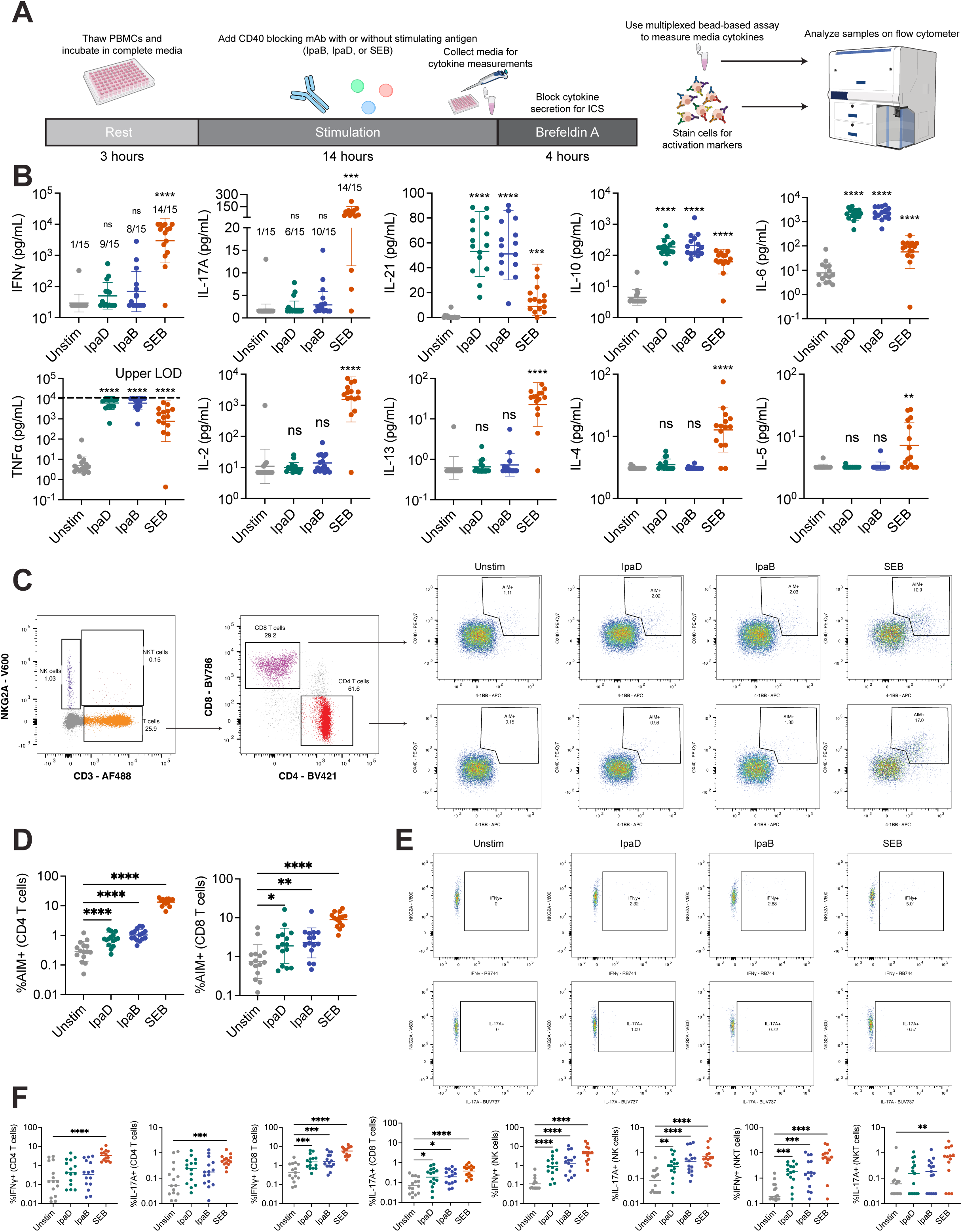
PBMC stimulation with IpaD and IpaB. (A) Schematic of experimental approach. Cryopreserved NHP PBMCs collected two-weeks post detection of infection were thawed, rested 3 h, and stimulated with IpaD (10 µg/mL), IpaB (10 µg/mL), or SEB (5 µg/mL) in the presence of anti-CD40 antibody. After 14 h, cytokines in media were quantified by multiplex assay. Brefeldin A was then added to block cytokine secretion. After 4 hours, the PBMCs were harvested and stained for flow cytometry analysis. (B) Concentrations of IFNγ, IL-17A, IL-21, IL-10, IL-6, TNFα, IL-2, IL-13, IL-4, and IL-5 detected in media collected following PBMC stimulations. Geometric mean ± geometric SD, one-way ANOVAs with Dunnett’s multiple comparisons tests. (C) Gating strategy for NK cells, NKT cells, CD4 T cells, CD8 T cells, and AIM^+^ (T cells OX40^+^ 4-1BB^+^). (D) Percent of total CD4 and CD8 T cells positive for AIM markers. Geometric mean ± geometric SD, one-way ANOVAs with Dunnett’s multiple comparisons tests. (E) Gating strategy for IFNγ^+^ and IL-17A^+^ CD4 T cells. (F) Percent of total CD4, CD8, NK, and NKT cells positive for IFNγ and IL-17A. Samples with 0%+ were assigned the lowest detected value. Geometric mean, one-way ANOVAs with Dunnett’s multiple comparisons tests. \**p*< 0.05, \*\**p<0.01*, \*\*\**p*< 0.001, and \*\*\*\**p*< 0.0001.

**Supplemental Figure 6:**
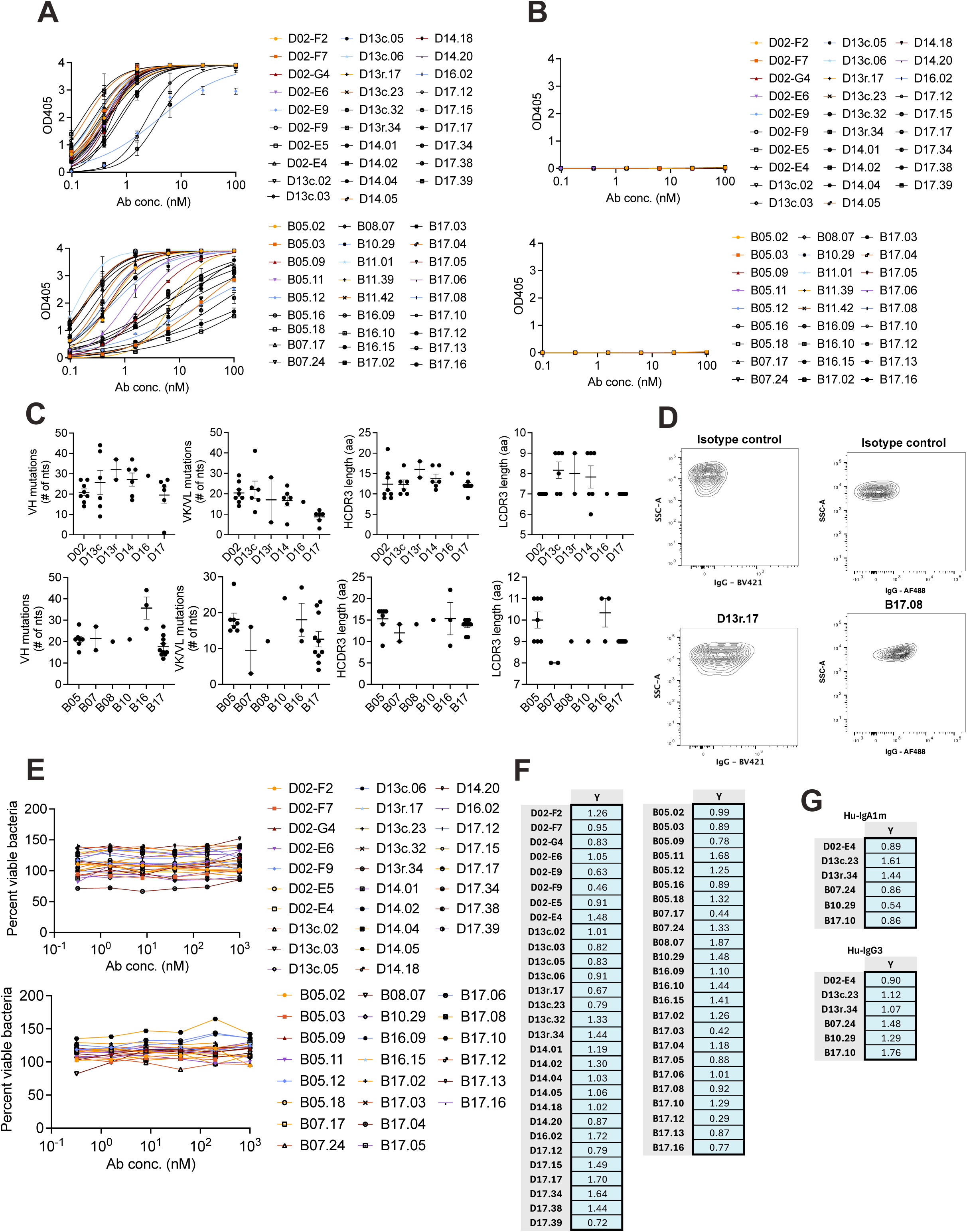
Characterization of T3SS mAbs, related to Figure 4. (A-B) ELISA curves of Ipa-specific mAbs against (A) IpaD or IpaB and (B) Ovalbumin. (C) Same data as Figure 4G. Mutation frequencies and CDR3 lengths of Ipa-specific mAbs by NHP. (n=1-8 mAbs per NHP) Ordinary one-way ANOVAs with Tukey’s post-test. (D) Representative flow plots of bacterial surface stains against *S. flexneri* 5a. (E) SBA curves for each Ipa-specific Mk-IgG1 mAb against *S. flexneri* Y. (F-G) Opsonophagocytosis assay with (F) Ipa-specific Mk-IgG1 mAbs or (G) isotype-switched Hu-IgA1 and Hu-IgG3 mAbs. Each value is the percent of all THP-1 cells positive for CellTrace Violet.

**Supplemental Figure 7:**
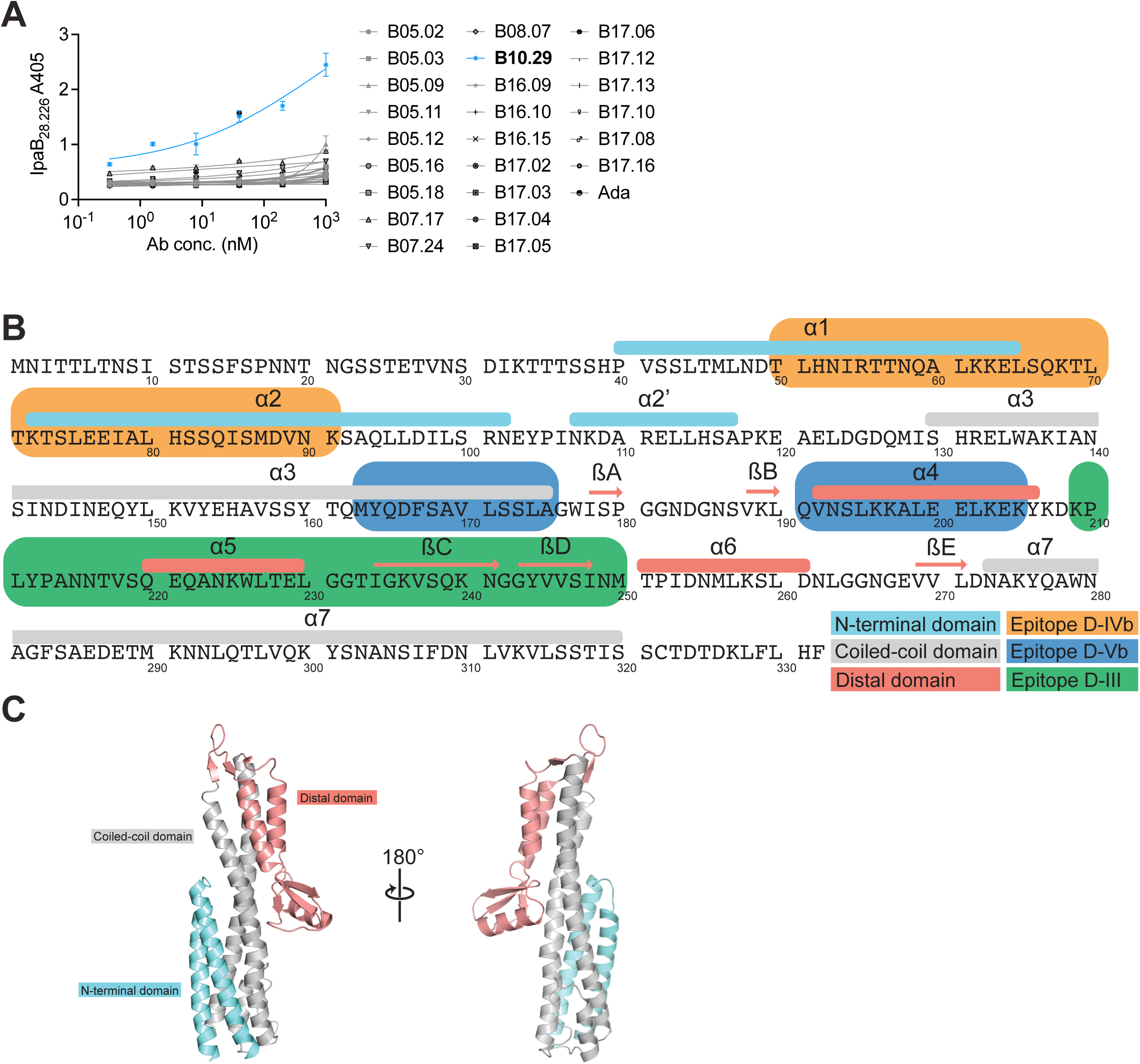
T3SS mAbs epitope mapping, related to Figures 5 & 6. (A) ELISA curves of IpaB-specific mAbs against IpaB_28.226_. Four parameter logistic curve fits. (B) Schematic depicting the primary and secondary structure of IpaD, with the approximate sites of epitopes D-IVb, D-Vb, and D-III, as well as the N-terminal, coiled-coil, and distal domains, indicated. (C) Model of an IpaD monomer (PDB 2J0O) with the N-terminal domain, coiled-coil domain, and distal domains colored blue, gray, and red, respectively.

**Supplemental Figure 8:**
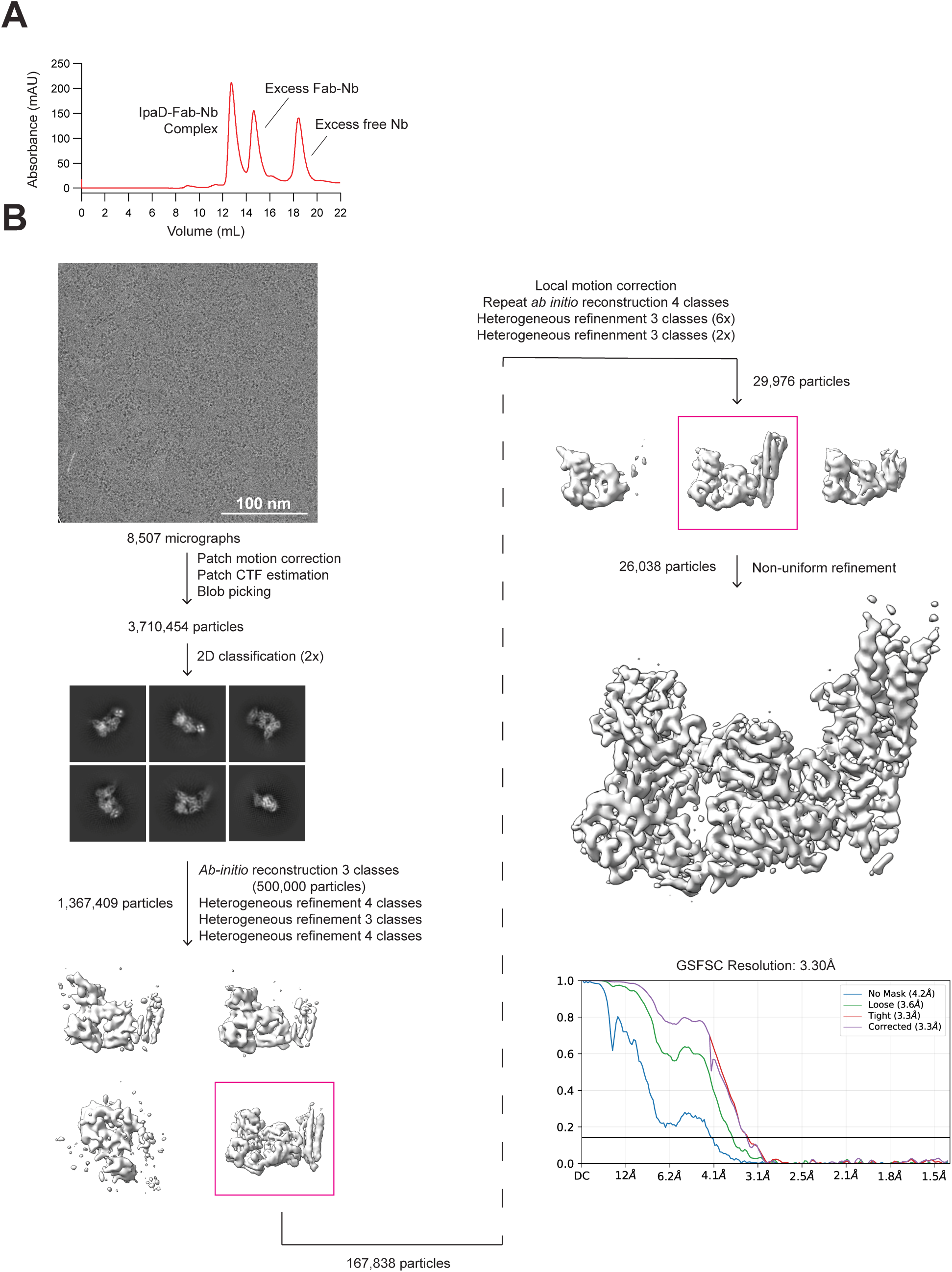
Cryo-EM data processing for the IpaD–D02-F2 Fab–D02-E4 Fab–anti-Fab Nb dataset, related to Figure 6. (A) Size-exclusion chromatography (SEC) trace for the IpaD–D02-F2 Fab–D02-E4 Fab–anti-Fab Nb complex shows clear separation of the complex from excess Fab-Nb complexes and excess free Nb. (B) Processing scheme used in CryoSPARC to determine the structure, including the final sharpened map and gold-standard Fourier shell correlation (FSC) curve calculated in cryoSPARC after FSC mask auto-tightening. The resolution was determined at FSC = 0.143 (horizontal black line). The final corrected mask gave an overall resolution of 3.4 Å.

**Supplemental Figure 9:**
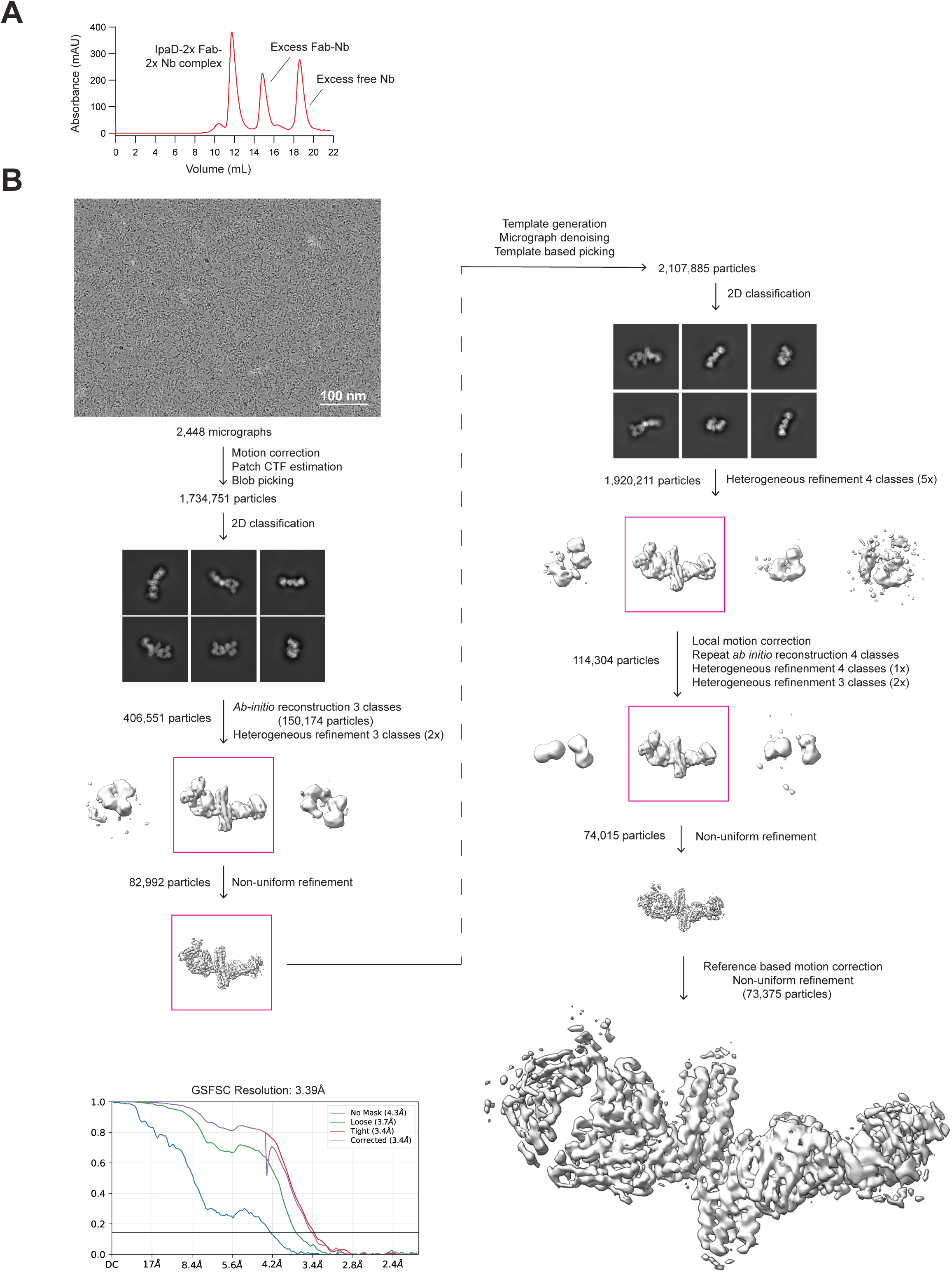
Cryo-EM data processing for the IpaD–D13-34 Fab–anti-Fab Nb dataset, related to Figure 6. (A) Size-exclusion chromatography (SEC) trace for the IpaD–D13-34 Fab–anti-Fab Nb complex shows clear separation of the complex from excess Fab-Nb complex and excess free Nb. (B) Processing scheme used in CryoSPARC to determine the complex structure, including the final sharpened map and gold-standard Fourier shell correlation (FSC) curve calculated in cryoSPARC after FSC mask auto-tightening. The resolution was determined at FSC = 0.143 (horizontal black line). The final corrected mask gave an overall resolution of 3.3 Å.

**Supplemental Figure 10:**
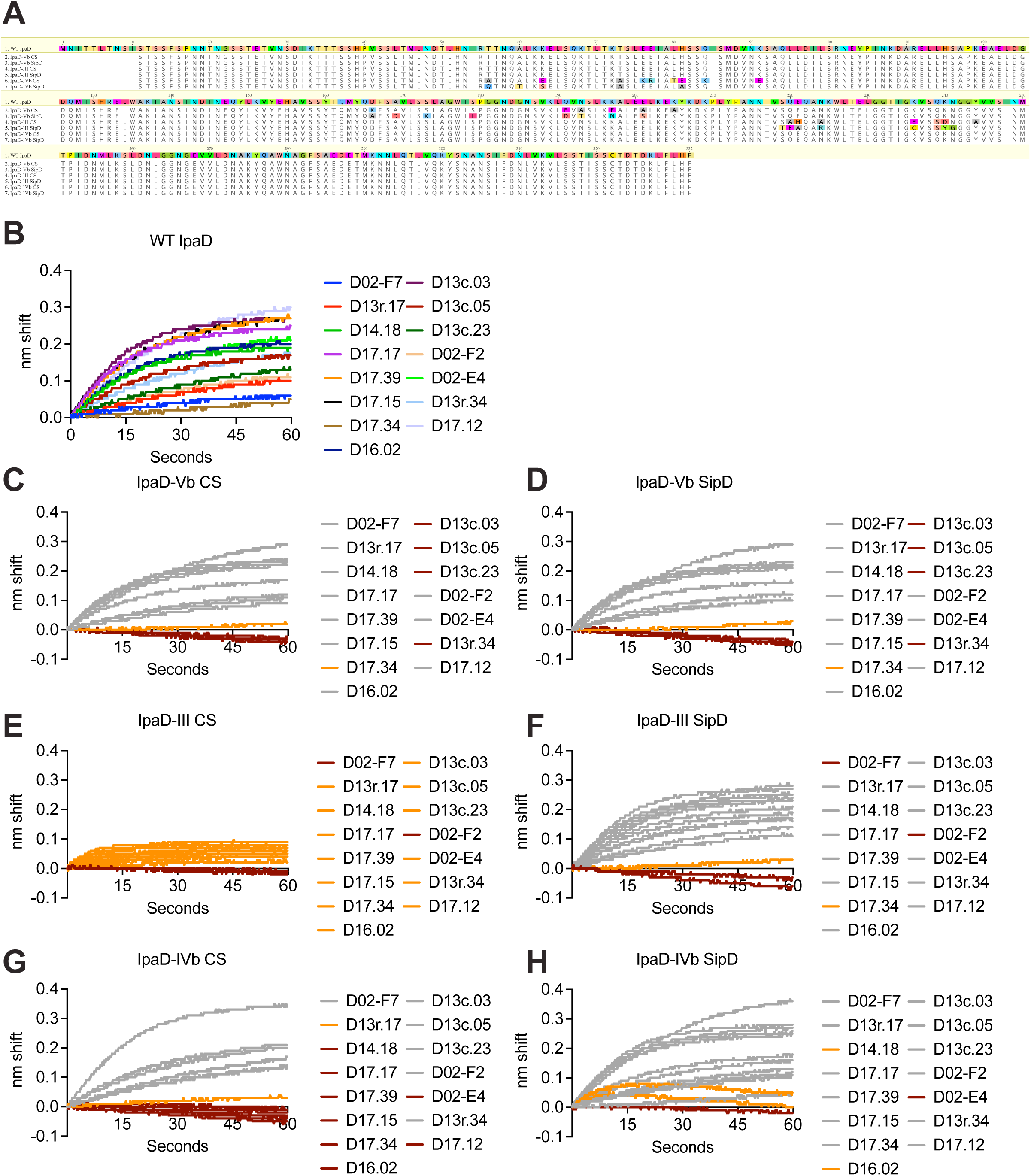
Epitope-specific IpaD mutagenesis, related to Figure 6. (A) Amino acid sequence of each mutagenized IpaD construct aligned to wild-type IpaD. (B-H) Biolayer interferometry traces of IpaD-specific antibodies binding to each IpaD construct loaded onto anti-HIS1K sensors.

## Methods

### NHP maintenance and regulatory approvals

Nonhuman primates were housed at the Wisconsin National Primate Research Center. Animal care and use were in accordance with the *Guide for the Care and Use of Laboratory Animals,*^87^ the Animal Welfare Act, and protocols approved by the College of Letters and Science and Vice

Chancellor for Research Institutional Animal Care and Use Committee of the University of Wisconsin-Madison. Animals were indoor housed on a 12-hour light/dark cycle with controlled temperature and humidity and were provided fresh water, commercially formulated monkey chow, fresh produce, and approved enrichment items daily. Animals were housed in groups, pairs, or singly when necessary, due to clinical, behavioral, or experimental reasons.

### Determination of NHP *Shigella* infection, bacterial isolation and characterization

NHP *Shigella* infections were determined by culture. Testing was performed on feces, rectal/fecal swabs, colon tissue, or gingival swabs collected from either individual NHPs or as pooled enclosure samples. Diagnostic specimens were collected and submitted for enteric bacterial culture to the UW Veterinary Care, Microbiology Laboratory. Specimens were inoculated to Trypticase soy agar containing 5% sheep blood, Hektoen Enteric agar (HE), and Selenite enrichment broth (Remel, Lenexa, KS) and incubated at 35 °C for 24 hours. The plates were examined for the presence of normal fecal microbiota and the HE agar was examined for typical non-lactose fermenting organisms resembling species of *Shigella*. The Selenite broth was subcultured to XLT4 media (Remel, Lenexa, KS) to screen for species of S*almonella.* Colonies resembling species of *Shigella* were subcultured to Trypticase soy broth containing 5% sheep blood and incubated for 24 hours at 35 °C. Identification and antimicrobial susceptibility testing were performed according to manufacturer recommendations (Vitek, bioMérieux, Durham, NC, USA) for isolates resembling species of *Shigella.* Antimicrobial susceptibility methods and breakpoint interpretations were in accordance with Clinical Laboratory Standards Institute guidelines^88^. Confirmed *Shigella* isolates were then swabbed and placed into liquid Amies medium for transport. Following transport, isolates were plated on tryptic soy agar with Congo red, and single red colonies were saved in glycerol stocks at -80 °C. Classical serotyping was performed using *Shigella* antisera (Subgroup B, Key Scientific Products). Overnight cultures of each isolate were stained with crystal violet in 20% methanol, then washed with DPBS. Suspensions of stained bacteria were gently mixed individually with the monovalent typing and grouping antisera and evaluated for agglutination and typed according to manufacturer’s key.

### Grading system for monitoring NHP symptoms

Animal demographics and health information were collected from WNPRC’s Electronic Health Records system. Symptoms described in these records were tracked from symptom onset to resolution corresponding to the culture sample collection date and were scored from 0-5 based on the severity of symptoms. A score of 0 represented asymptomatic cases. A score of 1-4 was assigned based on the number of observations present from the following list: abnormal feces, lethargy/down in cage/quiet, dehydration, vomiting, weight loss/inappetence, and pallor. A score of 5 represented cases where either euthanasia was elected or the animal was found dead. Abnormal feces were defined as reports of one or more of the following: bloody feces, mucoid feces, watery feces, or diarrhea.

### Bacterial strains and culture conditions

*Shigella* strains used for functional assays and OMV production in this study are listed in Table 1. For functional assays, frozen bacteria from glycerol stocks were streaked on TSBA plates with 0.01% Congo Red overnight at 37 °C (with 5 μg/mL kanamycin for strains harboring a kanamycin resistance cassette). Single colonies were then picked and grown shaking in TSB broth at 30 °C overnight and subcultured 1:100 the following day and grown shaking at 37 °C.

| Strain name | Species | Serotype | Origin |
| --- | --- | --- | --- |
| S022 | *flexneri* | Y | This study |
| 210059 | *flexneri* | 1b | Gift from Dr. Ferric Fang<sup>34</sup> |
| 2457T | *flexneri* | 2a | ATCC 700930 |
| CDC 3591-52 | *flexneri* | 2b | ATCC 12022 |
| J17B | *flexneri* | 3a | ATCC 24570 |
| S012 | *flexneri* | 3b | This study |
| M90T | *flexneri* | 5a | Gift from Dr. Eileen Barry<sup>89</sup> |
| CCH060 | *flexneri* | 6 | Gift from Dr. Eileen Barry |
| *S. flexneri* ΔwaaL (O-Ag knockout) | *flexneri* | N/A | Xerri et al.<sup>44</sup> |
| J17B *tolR::kanR* | *flexneri* | 3a | Xerri et al.<sup>44</sup> |
| 2457T *tolR::kanR* | *flexneri* | 2a | Xerri et al.<sup>44</sup> |
| 53G *Lvp::KanR* | *sonnei* | N/A | Provided by Baker lab |
| 53G *OAg::KanR* | *sonnei* | N/A | Provided by Baker lab |

### Bacterial genome sequencing and analysis

Bacteria were grown overnight in TSB (BD Biosciences), and then genomic DNA was extracted using Wizard® Genomic DNA Purification kit and quantified by Qubit™ dsDNA HS Assay kit. Genewiz (Azenta Life Sciences) conducted paired-end 150 bp Illumina sequencing on each sample. Galaxy USA suite of tools was used for genome analysis. Fastp^90^ was used to filter raw sequencing reads, then BWA-MEM2^91^ was used to map these reads to a reference genome.^35^ Bcftools consensus^92^ was then used to create consensus sequences for each isolate. Variants were then called using snippy (https://github.com/tseemann/snippy) and these data were used to construct a phylogenetic tree with IQ-TREE^93^. Antimicrobial resistance genes were identified using AMRFinderPlus^94^ in de novo assembled genomes constructed with Shovill (https://github.com/tseemann/shovill). The cladogram in Figure 1B was plotted using ggtree^95^ and the phylogenetic tree in Supplemental Figure 1A was plotted using iTOL.^96^

### NHP serum, plasma, and PBMC isolation

Blood was collected aseptically from NHPs into EDTA and serum-separating (SST) vacutainer tubes. Blood in the SST tubes was allowed to clot, then centrifuged at 4000 x g for 10 min. The serum supernatant was collected and centrifuged at 14,800 x g for 10 min at 4 °C to remove cell debris, then collected and heat inactivated at 56 °C for 30 min. Uncoagulated blood from EDTA tubes was pooled by sample and diluted with an equal volume of 1X DPBS. The diluted blood was carefully floated on top of prepared Lymphoprep (STEMCELL) in 50 mL SepMate tubes (STEMCELL) and centrifuged at 1200 x g for 20 min with the brake off. The plasma and lymphocyte layers were transferred to separate tubes and spun down at 650 x g for 8 min to pellet lymphocytes. Plasma was harvested and heat inactivated at 56 °C for 30 min. The pellet was resuspended in ACK Lysis Buffer (Gibco) for 5 min, then quenched with an equal volume of 1X DPBS and centrifuged at 550 x g. After discarding the supernatant, the pellet was washed in 5 mL of 1X DPBS and spun down at 400 x g. Cells were resuspended to approximately 2 x 10^7^ cells/mL in cold FBS. An equal volume of 20% DMSO in FBS was added dropwise to cell suspension before freezing and storing in LN_2_. Heat-inactivated plasma and serum were stored at -80 °C until thawing for use.

### Plasma cytokine determinations

Plasma cytokines were quantified by manufacturer’s instructions with the LEGENDplex^TM^ NHP Inflammation Panel. Data was collected using a BD FACSymphony and analyzed in FlowJo 10.10.0 (FlowJo LLC).

### Antigen preparation

IpaC was kindly provided as a gift by Dr. Marcela Pasetti from the University of Maryland Baltimore.

Production of IpaB has been described previously.^97,98^ Briefly, ipaB and *ipgC* (IpaB’s cognate chaperone) were cloned into pACYCDuet-1 with the IpaB being expressed with a His tag at the N-terminus and IpgC remaining untagged. The resulting plasmid was used to transform *E. coli* Tuner (DE3) for co-expression. The transformed bacteria were grown in LB containing chloramphenicol at 37 °C until A600 reached 0.8 at which protein expression was induced by addition of IPTG to 1 mM. After 3 h, bacteria were collected by centrifugation, washed and resuspended in IMAC binding buffer (20 mM Tris-HCl pH 7.9, 500 mM NaCl, 10 mM imidazole) with 0.1 mM AEBSF Protease Inhibitor and lysed using a microfluidizer at 18,000 psi with three passes. The cellular debris was removed by centrifugation at 10,000xg for 30 min and loaded onto a 5 ml HisTrap FF column. The IpaB/HT-IpgC was eluted with IMAC elution buffer (20 mM Tris-HCl pH 8.0, 500 mM NaCl, 500 mM imidazole), dialyzed into 50 mM Tris-HCl pH 8.0, and loaded onto a HiTrap Q FF column. The complex was eluted using a gradient of 50 mM Tris-HCl pH 8 containing 1M NaCl. Lauryldimethylamine oxide (LDAO) was then added to a final concentration of 0.1% to release the HT-IpgC. When the LDAO-treated L-DBF/HT-IpgC complex was passed over an IMAC column, the IpaB was collected in the flow-through with the HT-IpgC being retained on the IMAC column. Finally, IpaB was dialyzed into 20 mM phosphate, pH 7.2, with 150 mM NaCl (PBS) with 0.05% LDAO and stored at -80 °C. LPS levels were determined using a NexGen PTS with EndoSafe cartridges (Charles River Laboratories, Wilmington, MA). All proteins had LPS levels <5 Endotoxin units/mg.

IpaD, IpaB_28.226_, and mutagenized IpaD constructs (D13r.34 CS, D13r.34 SipD, D02-F2 CS, D02-F2 SipD, D02-E4 CS, and D02-E4 SipD) were cloned into a pET-28b(+) expression vector with N-terminal Avi- and/or His-tags. These plasmids were transformed into competent BL21 *E. coli* (NEB) by heat shock and selected on agar plates with kanamycin. Single colonies were picked, incubated overnight in LB-media supplemented with kanamycin, and subcultured 1:100 into fresh Terrific Broth (TB) supplemented with kanamycin. After reaching an OD_600_ of 0.6, 1 mM isopropyl-β-D-thiogalactopyranosid (IPTG) (Sigma) was added, and bacteria were incubated for an additional 18 h with shaking at 18 °C. Subsequently, bacteria were centrifuged at 4000 x g for 20 min, resuspended in resuspension buffer (20 mM Tris-HCl pH 8, 150 mM NaCl, 2 mM MgCl_2_, 1:10,000 benzonase, and cOmplete^TM^ EDTA-free protease inhibitor) then lysed with a probe sonicator. Recombinant protein was purified from the supernatant using HisPur^TM^ Ni-NTA Resin (Thermo Fisher Scientific) by gravity-flow column. IpaB_28.226_ was purified by anion exchange chromatography followed by size exchange chromatography, and mutagenized IpaD constructs were purified by size exchange chromatography. All protein yields were measured using NanoDrop A280, and purity was assessed by SDS-PAGE and size exclusion chromatography (SEC).

OMVs were purified as previously described in Xerri et al.^44^ Briefly, supernatant was collected from *S. flexneri* 2a or 3a tolR::kan mutants grown overnight in *Shigella sonnei* defined media (SSDM)^99^ supplemented with l-methionine and l-tryptophan, or an overnight culture of *S. flexneri* Y was sonicated 20 min in a bath sonicator at room temperature to induce OMV release. The supernatants were then filter-sterilized, concentrated by tangential flow filtration, and ultracentrifuged to pellet OMVs, which were then resuspended in PBS and stored at -20 °C. Total protein content of OMVs was quantified by Micro-BCA^TM^ assay (Thermo Fisher) following the manufacturer’s instructions.

O-Ag was purified from whole bacteria as previously described.^44^ In short, an overnight culture of *S. flexneri* Y was treated with acetic acid to hydrolyze the O-antigen. After hydrolysis, the culture was neutralized, bacterial debris was pelleted, the supernatant was buffer exchanged by tangential flow filtration (Sartorius) first into 1 M NaCl, and then into ddH2O. Citrate buffer (20 mM, pH 3) was added to the supernatant, incubated at RT for 30 min, debris was pelleted, and then the supernatant was passed through a Sartobind® IEX S75 cationic exchange filter (Sartorius). After cation exchange, the supernatant was precipitated once more with 18 mM Na2HPO4, 24% ethanol, and 200 mM CaCl_2_ at room temperature for 30 min, then buffer exchanged as described above before being concentrated in ddH2O using a 30 kDa cutoff spin filter (Amicon). The purified O-Ag was quantified using a Total Carbohydrate Assay Kit (Sigma) following the manufacturer’s instructions. Contaminating DNA and protein levels were assessed by nanodrop analysis and Micro BCA™ assay, respectively, and confirmed to be at least 10-fold lower than carbohydrate concentration.

### Antigen ELISAs

High-binding 96- or 384-well plates were coated with 5 µg/mL of protein antigen in PBS (IpaD, IpaB, or ovalbumin), or with 2.5 µg/mL of OMVs in PBS overnight at 4 °C. This solution was then discarded, and each well was blocked with 3% BSA, 0.1% Tween in PBS for protein antigens and 3% BSA, 0.01% Tween in PBS for OMVs at 37 °C for 1 h. Blocking solution was then discarded, and serial dilutions of serum or monkey IgG1 (Mk-IgG1) monoclonal antibody in PBS were added before incubating at 37 °C for 1 h. Plates were then washed 3x with 0.05% Tween in PBS for protein antigens and 0.001% Tween in PBS for OMVs. After washing with PBS-T, plates were incubated with AP-conjugated goat anti-monkey IgG antibody (Abcam) for 45 min at room temperature. After plates were washed again 3x with the appropriate PBS-T concentration, developing buffer (100 mM NaCl, 100 mM Tris-HCl pH 9.5, 50 mM MgCl_2_, 1% Tween 20, and 1 mg/mL phosphatase substrate) (Sigma) was added to each well. Plates were incubated for 20 min at room temperature before absorbance was measured at 405 nm using a BioTek Synergy multiplate reader.

### Serum Bactericidal Assay (SBA)

The desired *Shigella* strain was grown to mid-log phase in TSB (BD Biosciences), washed, and resuspended in SBA buffer (1X HBSS + 0.1% gelatin, pH 7.4).

In a 96-well V-bottom plate, 3x10^4^ CFU of *S. flexneri* were combined with 7.5% baby rabbit complement (BRC, Pel Freez Biologicals), the desired serum or monoclonal antibody, and SBA buffer in a final volume of 100µL. Plates were incubated at 37 °C for 3 h, washed once with 100 µL of SBA buffer, then mixed 1:1 with BacTiter-Glo^TM^ (Promega) in a white 96 well plate and incubated for 5 min at room temperature. Luminescence was detected on a BioTek Synergy plate reader. Each plate contained 6 wells of background buffer only controls and 6 wells of bacteria with no antibody controls. Background was subtracted from every well, then each well was normalized as a percentage of the bacteria with no antibody control wells. The data were fit with four parameter logistic curves with top values constrained to 100 and bottom values constrained to 0 in GraphPad Prism.

### Cell culture

THP-1 cells (ATCC, TIB-202) were cultured in RPMI complete media (RPMI 1640 Medium [Gibco], 4.5 g/L glucose, 1:100 GlutaMAX [Gibco], 10 mM HEPES [Gibco], 1 mM Sodium Pyruvate [Gibco], 55 µM beta-metcaptoethanol [Gibco], 10% fetal bovine serum [Omega Scientific], 50U/mL penicillin and 50 µg/mL streptomycin [Gibco]). Cells were passaged by transfer to fresh media without washing every 2-3 days and maintained at a density of 1 x 10^5^ – 5 x 10^5^ cells/mL. HeLa cells (ATCC, CCL-2) were passaged every 3-4 days in DMEM complete media (DMEM [Gibco], 1:100 GlutaMAX [Gibco], 10% fetal bovine serum [Omega Scientific], 50 U/mL penicillin and 50 µg/mL streptomycin [Gibco]). 0.25% Trypsin-EDTA (Gibco) was used to dissociate adherent cells from flask surface. Expi293F ^TM^ cells were passaged by transfer to fresh media without washing every 3-4 days in Expi293 ^TM^ Expression Medium (Gibco) and grown to a density of 3x10^6^ – 5x10^6^ cells/mL. Expi cells were cultured in shaker flasks using a rotating incubator. MS40L-low cells^100^ were passaged every 2-3 days in complete B cell media (RPMI 1640 Medium [Gibco], 1:100 GlutaMAX [Gibco], 10 mM HEPES [Gibco], 1 mM Sodium Pyruvate [Gibco], 1:100 MEM NEAA [Gibco], 55 µM beta-metcaptoethanol [Gibco], 10% fetal bovine serum [Omega Scientific], 50 U/mL penicillin and 50 µg/mL streptomycin [Gibco]).

### Opsonophagocytosis assay (OPA)

The desired *Shigella* strain was grown to mid-log phase in TSB (BD Biosciences), washed with PBS, then stained with a 1:50 dilution of CellTrace^TM^ Violet (CTV, Thermo Fisher) for 15 min at 37°C. Excess dye was quenched with FACS buffer (PBS + 1% FBS). The bacteria were washed once in PBS and resuspended in SBA buffer (1X HBSS + 0.1% gelatin, pH 7.4). 2.5 x 10^6^ CFU of bacteria were mixed with a 1:40 final dilution of either NHP serum or plasma, or 50nM of antibody and incubated at 37 °C for 30 min. Opsonized bacteria were then mixed with 5 x 10^4^ undifferentiated THP-1 cells (MOI of 50) and incubated for 30 min at 37 °C in 5% CO_2_. After incubation, the OPA reactions were fixed with 4% PFA in PBS on ice for 30 min. Cells were then washed in PBS and resuspended in FACS buffer (PBS + 1% FBS) before quantification by flow cytometry using a FACSCelesta or FACSymphony (BD Biosciences) and analysis via FlowJo 10.10.0 (FlowJo LLC).

### Adhesion and Invasion Assay (AIA)

One day before the assay, TC-treated 96-well plates were seeded with 4 x 10^4^ HeLa cells per well in 100 µL of DMEM complete media without antibiotics. The following day, *Shigella* was grown to mid-log phase in TSB (BD Biosciences), then stained with CTV as described in the OPA methodology. 20 µL of the bacteria resuspended to a concentration of 2.4 x 10^8^ CFU/mL was added to 96-well V-bottom plates pre-loaded with 5 µL of 1:10 dilutions of serum or plasma (final dilution of 1:50), then incubated at 37 °C for 30 min. 20 µL of this mixture was then transferred to each well of the prepared HeLa plate (MOI of 50) and centrifuged at 900xg for 10 min, then incubated at 37 °C for 30 min in 5% CO_2_. After incubation, cells were washed once with PBS, dissociated with Trypsin (Gibco), fixed with 100µL of 4% PFA in PBS, then incubated on ice for 30 mins. Cells were then centrifuged at 400 x g for 5 min and resuspended in FACS buffer (PBS + 1% FBS). Samples were then quantified by flow cytometry using a FACSCelesta or FACSymphony (BD Biosciences) and analyzed using FlowJo 10.10.0 (FlowJo LLC). % Inhibition was calculated relative to the invasion rate of bacteria pre-incubated in PBS.

### Hemolysis assay

The hemolysis assay was adapted from published studies.^101^ The desired *S. flexneri* strains were grown to mid-log phase in TSB (BD Biosciences). For assays with *S. sonnei*, an O-antigen knockout strain was used to expose the T3SS needle tip, and these cultures were supplemented with kanamycin. Cultures were centrifuged, washed in PBS, and resuspended in PBS to a final concentration of 6.6 CFU/mL. 45 µL of this suspension was added to each well of a 96-well V-bottom plate preloaded with 5 µL of 10 µM mAb or 5 µL of undiluted serum and incubated at 37°C for 30 min. Approximately 1 mL of sheep red blood cells (RBCs, Innovative Research) were washed 3 times with PBS and resuspended to a concentration of 1 x 10^9^ RBC/mL. 50 µL of washed RBCs was added to each well of the plate, centrifuged for 5 mins at 2000 x g, and incubated at 37 °C for 30 min. The hemolysis reactions were then stopped by adding 100µL of ice-cold PBS to each well and resuspending pellets with each addition. Plates were centrifuged again for 5 min at 2000 x g. 100 µL of supernatant was transferred from each well of the V-bottom plate to a new half-area 96 well plate. Plates were read at 545 nm on a BioTek Synergy multiplate reader. Each plate contained wells of RBCs with no bacteria to measure background and wells with bacteria and an isotype control antibody, Adalimumab,^102^ to measure baseline hemolysis. Background values were subtracted from every well, then each well was normalized as a percentage of hemolysis from the wells containing Adalimumab. Rabbit polyclonal IpaB antibody was obtained from Cusabio (CSB-PA322404ZA01SZB).

### NHP B cell sorts

Sorting of antigen-specific memory B cells was performed as previously described (citations, including that activation paper). Cryopreserved PBMCs were thawed and labeled with the following antibodies: CD3-APC-Cy7 (OKT3), CD4-APC-Cy7 (OKT4), CD8-APC-Cy7 (RPA-T8), CD14-APC-H7 (M5E2), CD16-APC-H7 (3G8), CD20-PerCP-Cy5.5 (2H7), IgM-BV785 (G20-127), and IgG-BV421 (G18-145), together with LIVE/DEAD Aqua Cell Stain in FACS buffer (PBS + 1% FBS). Cells were then stained with the antigen of interest (OMVs labeled with Vybrant^TM^ DiO or DiD solutions [Thermo Fisher], IpaD labeled with biotin [Avidity BirA500] and coupled to streptavidin-AF488 and -AF647 [Thermo Fisher], or IpaB labeled with NHS esters of AF488 and AF647 [Thermo Fisher]). Antigen specific memory B cells (CD3-, CD4-, CD8-, CD14-, CD16-, CD20+, IgM-, IgG+, antigen double positive) were sorted using a BD FACSMelody into individual wells of either an empty 96-well PCR plate or a 96-well cell culture plate pre-seeded with 3 x 10^3^ MS40-lo cells in 100 µL of complete B cell media supplemented with 5 µg/mL IL-2, 1 µg/mL IL-4, 1 µg/mL IL-21, and 1 µg/mL BAFF (PeproTech). PCR plates were immediately stored at -80 °C until PCR processing. Cell culture plates were maintained and passaged every 2-3 days to facilitate activation. 21 days post-sort, supernatants were assayed for antigen-specific binding by ELISA, and cells from positive wells were lysed in a buffer containing Igepal (Sigma), 0.1 M DTT and RNAseOUT (Thermo Fisher), transferred to a 96-well PCR plate, and stored at -80 °C until processing.

### Recombinant monoclonal antibody cloning and expression

Monoclonal IgG heavy and light chain sequences were amplified from either sorted single cells or activated cells frozen in lysis buffer that were stored at -80 °C. SuperScript ^TM^ IV Reverse Transcriptase and Buffer (Thermo) was used for RT-PCR in a master mix containing RNaseOUT ^TM^ (Thermo), random hexamer mix, 0.1 M DTT, dNTPs, and reverse primers for heavy, kappa, and lambda chains. The resulting cDNA was amplified via an initial PCR reaction (PCR1) using HotStarTaq MasterMix (Qiagen) and primers for heavy and light chains. A second PCR reaction (PCR2) was performed on samples to add Gibson overhangs at the beginning and end of amplified antibody chain sequences, which used a master mix containing Phusion ^TM^ Enzyme (Thermo), HF Buffer (Thermo), dNTPs, MgCl_2_, and primers for heavy and light chains. PCR2 products were screened via addition of SYBR Safe DNA stain (Thermo) and imaging in order to see which PCR wells successfully amplified a heavy, kappa, or lambda chain sequence. PCR2 hits were re-arrayed and run on a 1% agarose gel. After excising bands from gel, antibody chain sequences were isolated via ZR-96 Zymoclean Gel DNA Recovery Kit (Zymo) and assembled into Mk-IgG1 expression vectors via HiFi DNA Assembly (NEB). Assembled plasmids were transformed into NEB 5-alpha competent cells (NEB) and plasmid DNA was purified via either Zyppy-96 Plasmid Kit (Zymo) or NucleoSpin Plasmid Transfection-grade miniprep kit (Macherey-Nagel). Plasmid DNA was purified with Zyppy kit for initial colony screening via Sanger sequencing, while NucleoSpin kit was used to prepare final transfection-quality DNA. Expi293 ^TM^ Expression System was used for transfecting heavy and light chain antibody pairs. Transfections used a 1:2.5 heavy/light chain ratio by mass. FectoPRO ^TM^ transfection reagent (Avantor) was added to plasmid DNA diluted in Opti-MEM ^TM^ (Gibco) and incubated at room temperature for 10 min before Expi293F ^TM^ cells were added at a concentration of 3e6/mL. Transfections were left to express in rotating incubator for 5 days. 24 h post-transfection, cells were fed a mix of 300 mM Valproic acid sodium salt (Sigma) and 45% glucose solution (Sigma). Expressed monoclonal antibodies were purified 5 days post-transfection by centrifuging cells and incubating sterile-filtered supernatant with Praesto AP+50 beads (Purolite) for 3 h at 4 °C. Resin was transferred to Polyprep columns (Bio-Rad) and washed 3x with PBS by gravity-flow. IgGs were eluted using Pierce^TM^ IgG Elution Buffer (Thermo) and brought to a neutral pH with 1 M Tris-HCl pH 8.0 (Quality Biological). Antibody solutions were concentrated and buffer-exchanged into PBS using Amicon^TM^ Ultra-15 30 kDa filter units (Sigma).

### O-antigen and IpaD-specific mAbs bacterial surface stains

The desired *Shigella* strains were grown to mid-log phase in TSB (BD Biosciences). *S. flexneri* O-Ag-(to negatively screen for O-Ag-specific mAbs), *S. sonnei* lvp::kan (to stabilize the virulence plasmid for O-Ag-specific mAbs staining), and *S. sonnei* OAg::kan (to remove the O-Ag and expose the T3SS needle for Ipa mAbs staining) were cultured in TSB supplemented with kanamycin.

For IpaD and OAg-specific antibodies, bacteria were pelleted, washed, and resuspended in PBS to a concentration of 2 x 10^9^ CFU/mL. In a 96-well V-bottom plate, 25 µL of bacteria were added to 25 µL of antibody at 2X final concentration and incubated at 37°C for 30 mins. The bacteria were then washed in PBS and resuspended in 25 µL of secondary antibody (BD Horizon™ BV421 Mouse Anti-Human IgG), diluted 1:100 in PBS and incubated for 30 min at room temperature. To fix the bacteria, 100 µL of 4% PFA in PBS was added to each well and the plates were incubated on ice for at least 30 min. Bacteria were pelleted and then resuspended in FACS buffer (PBS + 1% FBS). Fluorescence was read on a BD FACSCelesta or BD FACSymphony. Data was analyzed on FlowJo 10.10.0 (FlowJo LLC).

### IpaB-specific mAbs bacterial surface stains

*Shigella* strains were grown to mid-log phase in fresh TSB (BD Biosciences) supplemented with 0.1% deoxycholate (Sigma). In a 96-well V-bottom plate, 25 µL of unwashed bacterial culture was added to 25 µL of pre-warmed antibody at 2X final concentration, working quickly to avoid the culture cooling, and incubated at 37 °C for 30 min. 100 µL of ice cold 4% PFA in PBS was then directly added to the plate and incubated on ice for 15 min. Bacteria were washed with PBS and resuspended in 1:100 unconjugated mouse anti-human IgG antibody (G18-145, BD Biosciences) in PBS followed by a 30 min incubation at 37 °C. Bacteria were washed with PBS and resuspended in 1:100 AF488-conjugated goat anti-mouse IgG (Thermo Fisher) followed by another 30 min incubation at 37 °C. Bacteria were washed with PBS, resuspended in FACS buffer, and quantified on a BD FACSymphony. Data was analyzed on FlowJo 10.10.0 (FlowJo LLC).

### Antibody isotype switching / purifications

Heavy and light chain sequences from representative mAbs were PCR-amplified from IgG1 plasmid DNA using primers with proper overhang sequences following the same Phusion ^TM^ protocol as our PCR2 cloning step. PCR products were run on a 1% agarose gel and gel extracted via ZR-96 Zymoclean Gel DNA Recovery Kit (Zymo). Heavy and light chain pairs were Gibson assembled into parent vectors to express antibodies in the form of IgG3 and monomeric IgA1 via HiFi DNA Assembly (NEB). Plasmid DNA was purified using Zyppy-96 Plasmid Kit (Zymo) and screened for correct assembly via Sanger sequencing. Sequenced plasmids were purified via NucleoSpin Plasmid Transfection-grade miniprep kit (Macherey-Nagel). Heavy and light chain pairs were transfected following the same transfection protocol as previously described, with the exception of monomeric IgA1 using a 1:1 heavy/light chain ratio. 5 days post-transfection, cells were centrifuged and supernatants were sterile-filtered and incubated for 3 h at 4 °C with either Protein A/G UltraLink ^TM^ Resin (Thermo) for IgG3s or Peptide M Agarose Resin (Invivogen) for IgA1s. Antibodies were purified and buffer-exchanged following the same column purification procedure as previously described. Purified IgG3 and IgA1 antibodies were run on a Mini-PROTEAN TGX 4-20% Precast Gel for size validation (Bio-Rad).

### Mouse work

Mouse experiments were approved by the Office of Animal Research (OAR) and the institutional IACUC at the University of Missouri (Protocol: 38241). Six-to-eight-week-old female C57BL/6 mice (Charles River Laboratories, Wilmington, MA) were used in this study. Groups were as follows (n=10): isotypic control mAb VRC01 (negative control); mAb D13r.34, D02-E4, B07.24, B10.29. Mice received a single intranasal dose of antibody at 10 mg/kg in a total volume of 30 – 50 µL (based on the weight of each mouse). Antibody administration was performed 6 h prior to Shigella challenge. S. flexneri 2a 2457T challenge strains were grown on tryptic soy agar containing 0.025% Congo red at 37 °C overnight, subcultured in tryptic soy broth on the challenge day to an A600 of 1.0, and diluted to 5 x 10^5^ CFU (LD50) in 30 µl PBS for intranasal (IN) challenge. Mice were monitored for two weeks, with euthanasia criteria including >25% weight loss or blood glucose ≤100 mg/dL. This experiment was conducted two independent times, and data was combined for analysis.

### PBMC stimulation assays

Cryopreserved PBMCs were thawed and seeded into TC-treated 96 well plates at a density of 7 x 10^5^ cells per well in RPMI complete media (RPMI 1640 Medium [Gibco], 4.5 g/L glucose, 1:100 GlutaMAX [Gibco], 10 mM HEPES [Gibco], 1 mM Sodium Pyruvate [Gibco], 55 µM beta-metcaptoethanol [Gibco], 10% fetal bovine serum [Omega Scientific], 50 U/mL penicillin and 50µg/mL streptomycin [Gibco]). The cells were rested for 3 h at 37 °C with 90% humidity and 5% CO_2_. CD40 blocking antibody (1 µg/mL final conc.) and stimulating antigen (Ipa proteins 10 µg/mL, SEB 5µg/mL final conc.) were added to each well, bringing total volume per well to 200 µL. After 14 h of incubation, 100 µL of media was collected for cytokine determinations using the LEGENDplex^TM^ NHP Th Cytokine Panel (BioLegend) according to the manufacturer’s instructions. 10µL of Brefeldin pre-diluted 1:100 in RPMI complete media (final dilution 1:1,000) was then added. After 4 h more of incubation, cells were dissociated with Trypsin (Gibco) and transferred to 96-well V-bottom plates. Cells were stained in FACS buffer with the following antibodies: CD3-AF488 (SP34-2), CD4-BV421 (OKT4), CD8-APC-Cy7 (RPA-T8), NKG2A-Vio Bright V600 (REA110), OX40-PE-Cy7 (Ber-ACT35), and 4-1BB-APC (4B4-1), along with LIVE/DEAD Near-IR Cell Stain. Cells were then fixed in BD CytoFix, permeabilized in BD CytoPerm, and stained for intracellular cytokines with the antibodies IFNγ-RB744 (B27), and IL-17A-BUV737 (N49-653). Cells were then resuspended in FACS buffer and quantified on a BD FACSymphony, then the data was analyzed using FlowJo 10.10.0 (FlowJo LLC).

### Competition ELISA

384-well plates were coated with 5 µg/mL of each antibody in PBS to be assayed in duplicate, then incubated overnight at 4 °C. This liquid was then discarded, and plates were blocked with 3% BSA, 0.1% Tween in PBS at 37 °C for 1 h. While the plates were blocking, 150 µg/mL of competing mAb was pre-incubated with either 1 µg/mL IpaD in PBS or 60 µg/mL IpaB in TBS+LDAO 0.05% at 37 °C for 30 min. The blocking solution was then discarded, and the antigen-antibody complexes were then transferred to the blocked plates and incubated at 37 °C for 1 h. The plates were then washed 3 times with either PBS+0.05% Tween for IpaD ELISAs or TBS+0.05% LDAO for IpaB ELISAs. 1:1000 of AP-conjugated mouse anti-His antibody (SB194b, SouthernBiotech) diluted in either PBS for IpaD or TBS+0.05% LDAO for IpaB was added to each well and incubated at 37 °C for 30 min. The plates were again washed three times with the appropriate buffer, then AP developing buffer was added to each well and incubated for 25 min at room temperature. Absorbance at 405 nm was measured using a BioTek Synergy multiplate reader. Self-competing wells were subtracted from each antibody as background values, then each antibody combination was normalized as a percentage of the signal from the unblocked coating antibody. These data were then used to create a Pearson correlation matrix and plotted in heatmap form.

### Cryo-EM sample preparation and data collection

All IpaD Fabs were prepared by papain digest. The IpaD–D02-F2–D02-E4 complex was generated by combining purified IpaD with a 1.5-fold molar excess of D02-F2 Fab, 1.5-fold molar excess of D02-E4 Fab, and 6-fold molar excess of a previously described anti-Fab nanobody (Nb).^103^ The complex was isolated using a Superdex 200 Increase (Cytiva) SEC column equilibrated in HEPES-buffered saline (HBS). The final complex was concentrated to an absorbance at 280 nm (A280) of 0.57. The IpaD–D13r.34 complex was generated similarly, except that a 1.5-fold molar excess of D13-34 Fab and 3-fold excess of the anti-Fab nanobody were incubated with IpaD prior to SEC, and the final sample was concentrated to an A280 of 0.4. UltrAuFoil grids with 1.2-μm diameter/1.3-μm spacing (quantifoil) were plasma cleaned for 30 sec. Grids were prepared using a Vitrobot Mark IV (ThermoFisher) set to 6 °C and 100% humidity using 4 µL sample with a 10 s wait time and a 4 s blot time at force 15 (IpaD–D02-F2–D02-E4) or force 12 (IpaD–D13-34) before plunging into liquid ethane. Data for the IpaD–D02-F2–D02-E4 sample were collected on a Talos Arctica operating at 200 kV (ThermoFisher Scientific) equipped with a K3 direct electron detector (Gatan) using Serial EM. For the IpaD–D13-34 sample, data were collected on a Titan Krios operated at 300 kV (ThermoFisher Scientific) equipped with a Falcon4i direct electron detector and a Selectris Energy Filter using EPU. See Table 1 for data collection parameters.

### Cryo-EM data processing and model building

For the IpaD–D02-F2–D02-E4 dataset, motion correction and dose weighting, followed by patch contrast transfer function (CTF) estimation, blob picking, and patch motion correction were performed in CryoSPARC Live.^104,105^ A stack of 1,734,751 particles was picked using CryoSPARC Live Blob Picker. Particles were subjected to one round of 2D classification, *ab initio* reconstruction, and three rounds of 3D heterogeneous refinement that resulted in a stack of 82,992 particles. Non-uniform refinement^106^ was used to generate a map that was subsequently used for template-based particle picking on de-noised micrographs. A total of 2,107,885 particles were picked and extracted with 2x Fourier cropping. After multiple rounds of 2D classification and 3D heterogeneous refinement, a particle stack of 114,304 was subjected to local motion correction and extracted without binning. Heterogenous refinement was repeated until the map quality no longer improved, leaving a particle stack of 74,015, which was reduced to 73,375 during the final round of non-uniform refinement with reference-based motion correction.^105^

For the IpaD–D13r.34 dataset, patch motion correction and dose weighting, followed by patch contrast transfer function (CTF) estimation was performed in CryoSPARC. Blob picker was used to generate an initial stack of 3,710,545 particles which were extracted with 2x Fourier cropping. Two rounds of 2D classification were used to reduce the particle stack to 1,367,409. A subset of 500,000 particles was used to generate four *Ab initio* models. After three rounds of 3D heterogeneous refinement, a stack of 167,838 particles was subjected to local motion correction and extraction without binning. Heterogenous refinement was repeated until the map quality no longer improved, leaving a particle stack of 26,038. Local CTF refinement and reference-based motion correction did not improve map quality. Masking the constant region of the antibody fragment and the anti-Fab nanobody did not improve the quality of the final map in either dataset, which was highest when using automatic mask generation in non-uniform refinement.

The coordinates were built manually in Coot and ISOLDE,^107,108^ using starting models generated from AlphaFold3.^109^ The anti-Fab nanobody and the constant region of the antibody fragments were not included in the models. The structures were refined using real-space refinement in Phenix.^110,111^ Structural biology software used in this project was compiled and configured by SBGrid.^112^ Figures were generated using PyMOL and ChimeraX.^113^

### Biolayer interferometry (BLI) assays

BLI assays were performed using polypropylene black 96-well microplates (Greiner) on an Octet R8 (Sartorius) instrument at 30 °C. All reagents were diluted with 10X kinetics buffer (0.1% BSA, 0.02% Tween 20 in 1XPBS). 8 µM of His-tagged protein was captured with HIS1K Biosensors (Sartorius) for 60 seconds and transferred to 10X kinetics buffer for 30 seconds to wash off unbound protein. Sensors were then transferred into 30nM of monoclonal antibodies for 60 seconds. The percent WT shift for each mAb was calculated with the formula: Percent WT shift (%) = (IpaD construct shift @ 60s [nm]) / (WT IpaD shift @ 60s [nm]) * 100.

